# Comparative genome analysis of *Enterococcus cecorum* reveals intercontinental spread of a lineage of clinical poultry isolates

**DOI:** 10.1101/2022.10.18.512807

**Authors:** Jeanne Laurentie, Valentin Loux, Christelle Hennequet-Antier, Emilie Chambellon, Julien Deschamps, Angélina Trotereau, Sylviane Furlan, Claire Darrigo, Florent Kempf, Julie Lao, Marine Milhes, Céline Roques, Benoit Quinquis, Céline Vandecasteele, Roxane Boyer, Olivier Bouchez, Francis Repoila, Jean Le Guennec, Hélène Chiapello, Romain Briandet, Emmanuelle Helloin, Catherine Schouler, Isabelle Kempf, Pascale Serror

## Abstract

*Enterococcus cecorum* is an emerging pathogen responsible for osteomyelitis, spondylitis, and femoral head necrosis causing animal suffering, mortality, and requiring antimicrobial use in poultry. Paradoxically, *E. cecorum* is a common inhabitant of the intestinal microbiota of adult chickens. Despite evidence suggesting the existence of clones with pathogenic potential, the genetic and phenotypic relatedness of disease-associated isolates remains little investigated. Here, we sequenced and analyzed the genomes and characterized the phenotypes of more than 100 isolates, the majority of which were collected over the last ten years in 16 French broiler farms. Comparative genomics, genome-wide association study, and measured susceptibility to serum, biofilm forming capacity, and adhesion to chicken type II collagen were used to identify features associated with clinical isolates. We found that none of the tested phenotypes could discriminate origin of the isolates or phylogenetic group. Instead, we found that most clinical isolates are grouped phylogenetically and our analyses selected six genes that discriminate 94% of isolates associated with disease from those that are not. Analysis of the resistome and the mobilome revealed that multidrug-resistant clones of *E. cecorum* cluster in few clades and that integrative conjugative elements and genomic islands are the main carriers of antimicrobial resistance. This comprehensive genomic analysis shows that disease-associated clones of *E. cecorum* belong mainly to one phylogenetic clade.

**IMPORTANCE:** *Enterococcus cecorum* is an important pathogen in poultry worldwide. It causes a number of locomotor disorders and septicemia, particularly in fast-growing broilers. Animal suffering, antimicrobial use, and associated economic losses require a better understanding of disease-associated *E. cecorum* isolates. To address this need, we performed whole genome sequencing and analysis of a large collection of isolates responsible for outbreaks in France. By providing the first dataset on the genetic diversity and resistome of *E. cecorum* strains circulating in France, we pinpoint an epidemic lineage probably also circulating elsewhere and which should be targeted preferentially by preventive strategies in order to reduce the burden of *E. cecorum*-related diseases.

## INTRODUCTION

*Enterococcus cecorum* is a commensal bacterium of the gut microbiota of adult chicken (1–3). This bacterium has emerged over the last twenty years as a significant cause of locomotor disorders in poultry worldwide, particularly in fast-growing broilers (4). In France, reports on *E. cecorum* between 2006 and 2018 have shown an increase from 0.3% to more than 7% of total avian infections, the majority of which include locomotor disorders in broilers (5). *E. cecorum* is mostly responsible for osteomyelitis, spondylitis, vertebral osteoarthritis, and femoral head necrosis (1, 6, 7), causing substantial losses in broiler production due to culling, mortality, condemnations at the slaughterhouse, veterinary costs, and increased exposure to antibiotics (6, 8, 9). The development of *E. cecorum* infections is multifactorial and depends on host genetics, rapid growth, feed composition, husbandry procedures, and animal density in combination with the pathogenic potential of the bacterium (10–13). The association between early intestinal carriage of *E. cecorum* and increased risk of infections points to the gastro-intestinal tract as a route of infection (10, 11, 14, 15). Recent studies indicate that lesion-associated *E. cecorum* isolates appear to be adapted to colonize the gut early in life, in contrast to non-clinical isolates (i.e., strains isolated from the gut of healthy birds) that do not colonize to a detectable level before week 3 (10, 14). It was also proposed that disinfection failure may contribute to *E. cecorum* persistence and outbreaks due to biofilm formation (7, 16–18). Although suggested by the prediction of host-binding proteins in the genome (19), the adhesion to host tissue proteins has been overlooked and the robustness of *E. cecorum* biofilms and associated properties remain to be investigated. It is also assumed that in Gram-positive bacteria, the thick layer of peptidoglycan that surrounds the cytoplasmic membrane confers resistance to the bactericidal activity of the serum: for instance, human serum selectively kills commensal *Enterococcus faecium* strains whereas disease-associated *E. faecium* strains are not susceptible (20). Assessing the pathogenic potential of *E. cecorum* isolates remains a challenge. Recently, an *in vivo* model has been used to distinguish the pathogenicity between two clinical isolates under field conditions, but it is applicable to only a limited number of strains (3). Less limiting, the chicken embryo lethality assay (CELA) has shown tendencies where pathogenic strains kill more efficiently than commensal isolates (14, 21).

Several molecular epidemiological studies based on pulse-field gel electrophoresis (PFGE) patterns of commensal and clinical isolates from the United States, Canada, Belgium, the Netherlands, Germany, and Poland agree that commensal isolates have a higher diversity than clinical isolates, suggesting the evolution of specific clones with higher pathogenic potential. However, clinical isolates exhibited multiple PFGE patterns, supporting the hypothesis of the polyclonal nature of the infectious isolates (22–26). Furthermore, repeated outbreaks with genotypically related isolates within farms and local areas substantiate horizontal transmission and a farm-related reservoir (7, 17, 24, 26, 27). To date, only one complete genome of *E. cecorum* is available (type strain NCTC-12421 accession number NZ_LS483306.1) and only two comparative genomic studies of *E. cecorum* isolates from the United States have been performed (19, 28). Comparison of partial genomes of three commensal and three clinical isolates from the southeastern United States isolated between 2010 and 2011 indicated that the pathogenic *E. cecorum* strains had smaller genomes with more than 120 genes absent or whose products had less than 40% identity in the commensal isolates (19). On the other hand, ~70 genes of the non-clinical isolates were absent or encoded products with less than 60% identity in the clinical isolates. In line with studies reporting a high rate of clinical isolates unable to metabolize mannitol (14, 24, 25), the orthologs of mannitol phosphate dehydrogenase, the mannitol operon activator, as well as the mannitol-specific component IIA of the phosphotransferase system (PTS) were not found in clinical isolates. In another other study, partial genomes of nine clinical isolates isolated in Pennsylvania in 2008 and 2009 were compared with those of nine non-clinical isolates from the National Antimicrobial Resistance Monitoring System isolated between 2003 and 2010 (28). The trend of a slightly smaller genome size for clinical isolates was confirmed and consistent with a larger accessory genome of non-clinical isolates. Noticeably, the non-clinical genomes had more antibiotic resistance genes. By combining available *E. cecorum* draft genome sequences (29, 30), the core genome was estimated to be 1,436 genes (28). Phylogenetic analysis of the core genome led the authors to conclude that the isolates cluster independently of their clinical or non-clinical status, which raises the question of whether the clinical isolates of *E. cecorum* belong to specific genetic groups. The objective of this study was to provide a better insight on genomic organization and phenotypic diversity of *E. cecorum* clinical isolates from broilers circulating between 2007 and 2017 in Brittany, the leading French commercial broiler producing area. We performed whole genome sequencing of more than hundred poultry and human clinical isolates in order to better define the extent of genetic relatedness of clinical isolates and detect genes associated with virulence-related traits. We completed this genomic analysis by testing isolates for their adhesion to type II collagen, biofilm robustness, and growth in chicken serum. The overall genetic diversity of *E. cecorum* was investigated by pan-genome analysis, with a particular focus on mobile genetic elements (MGEs), antimicrobial resistance genes (ARGs), and genome-wide associations (GWAS) between the accessory genes and the phenotypic traits.

## RESULTS

### *Clonality of* E. cecorum *clinical isolates*

To get insight into the gene repertoire of *E. cecorum*, we performed whole genome sequencing of 118 isolates, including 100 clinical isolates collected from 16 broiler farms in western France between 2007 and 2017, 6 clinical isolates of human origin, and 12 isolates from other studies that had been previously sequenced (Table S1). 118 genomes sequenced by Illumina technology had a sequencing coverage greater than 160X and an N50 between 62 kbp and 276 kbp (Table S2A). Hybrid assembly of Illumina and Nanopore data of 14 genomes allowed the reconstruction of 10 complete genomes (CIRMBP-1212, CIRMBP-1228, CIRMBP-1246, CIRMBP-1261, CIRMBP-1274, CIRMBP-1281, CIRMBP-1283, CIRMBP-1287, CIRMBP-1292, and CIRMBP-1302) and improvement of genome assembly completeness of 4 others (Table S2A). The estimated average length of the genomes is ~2.4 Mb and varies between ~2.05 and ~2.8 Mbp. Each genome had an average of 2,345 predicted protein coding sequences (CDSs). A total of 277,011 CDSs were annotated. Comparison of the chromosomal architecture of the 11 complete *E. cecorum* strains using NCTC12421 strain as reference revealed that strain CIRMBP-1261 has a large chromosomal inversion of ~1.8 Mbp between the second and sixth ribosomal RNA operon (Fig. S1). Strain CIRMBP-1287 also has a chromosomal inversion of 280 kbp from genes DQL78_RS05120 to DQL78_RS06585, involving an insertion sequence (IS) of the IS3 family.

A further thirty available non-redundant *E. cecorum* genomes were included. These comprised genomes of 9 clinical and 21 non-clinical isolates from Belgium, Germany, and the United States (Table S2B). Comparative genomics analysis of the 351,733 CDSs from the 148 genomes identified 8,523 gene clusters in the pan-genome composed of a strict core-genome (present in all *E. cecorum* genomes) of 1,207 CDSs, an accessory genome of 4,664 CDSs, and a unique genome of 2,652 (31.1%) CDSs (Fig. 1). The pan-genome curve displayed an asymptotic trend after the 140-genome iteration, indicating a stabilization of the pan-genome within this dataset. Consistently, the core-genome stabilized after the 125-genome iteration (Fig. 1A). These trends confirm that the genome dataset used here provides a comprehensive overview of the gene repertoire of the *E. cecorum* species. The distribution of the gene clusters revealed that ~47% of the pan-genome (n=3,979 including the unique genes) are present in one to three isolates (Fig. 1B), indicating that the genetic diversity is partly attributable to gene acquisition. This hypothesis is further supported by a high proportion (42%) of genes of unknown function.

**Fig. 1:**
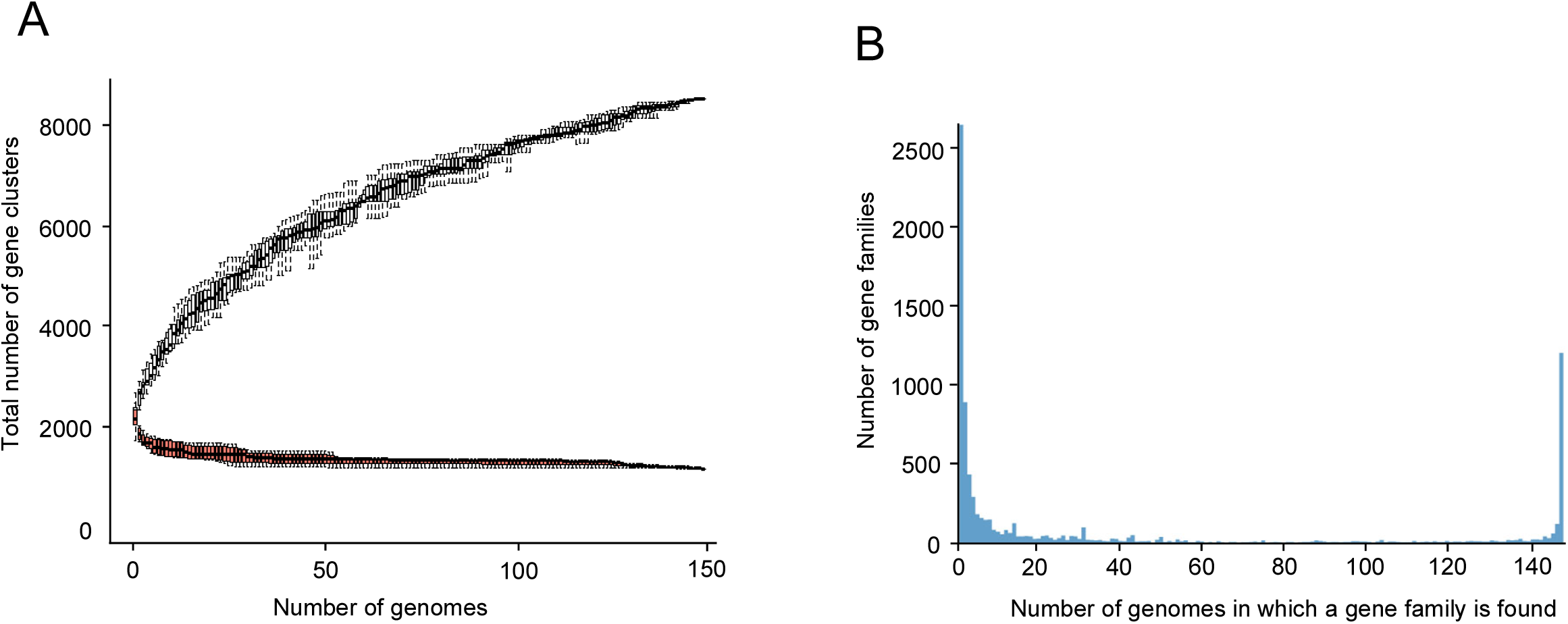
Pan-genome analysis of 148 *E. cecorum* genomes. **A.** Accumulation curves of gene clusters of the pan and core-genomes. Pan-genome size in black corresponds to the total number of gene clusters against the number of genomes included. Core-genome size in red corresponds to the number of gene clusters in common against the number of genomes included; numbers averaged on 1,000 randomized orders for genome addition. **B.** Gene frequency spectrum. Only one representative per gene cluster is considered.

The *E. cecorum* neighbor-joining (BioNJ) phylogenetic tree was constructed using the concatenated sequence of the core genes (Fig. 2). The isolates clustered in five distinct phylogenetic clades (A to E), supported by a bootstrap value of 100% and a higher maximum pairwise genetic distance between clades (0.026 to 0.041) than within clades (<0.021) (Fig. S2). Clade E was further divided into 13 well-supported subclades (E1 to E13) with intra subclade pairwise genetic distances below 0.012, indicating higher clonality of these isolates. Clades A to D contain 2 to 16 genomes, of which only 35% were avian clinical isolates. While clades B and D contain mainly genomes of non-clinical isolates from the United States, clade A contains two European clinical human isolates and clade C contains both non-clinical and clinical poultry isolates. In contrast, clade E comprises 117 genomes, 95% of which belong to avian clinical isolates from the United States, Belgium, Germany, Poland, and France, suggesting a widespread-distribution of this clade. Of note, the type strain NCTC12421, isolated from caecal content of a dead chicken from a farm in Belgium (1, 31) is part of subclade E3 and subclade E12 contains only avian isolates from the United States. Although the number of isolates per subclade is limited, almost all isolates from subclades E6, E10, and E13 were isolated after 2009 while those from subclades E4 and E11 were isolated before 2014 and after 2015, respectively, indicating that the dominant subclasses have varied over the years. Due to sequence data availability, single nucleotide polymorphism (SNP) analysis was only possible for genomes sequenced in this study, thus excluding genomes from clades B, D and subclade E12. Of the 65,226 SNPs of the core-genome 2,443, 168, and 62 were specific to isolates of clades A (n=2), C (n=9) and E (n=107), respectively. Of the thirteen non-synonymous clade E-specific SNPs, eight are non-conservative and two are predicted as nonneutral in a phage shock protein (PspC, DQL78_RS04285 in NCTC12421) and ATP-binding cassette domain-containing protein (DQL78_RS04370 in NCTC12421).

**Fig. 2:**
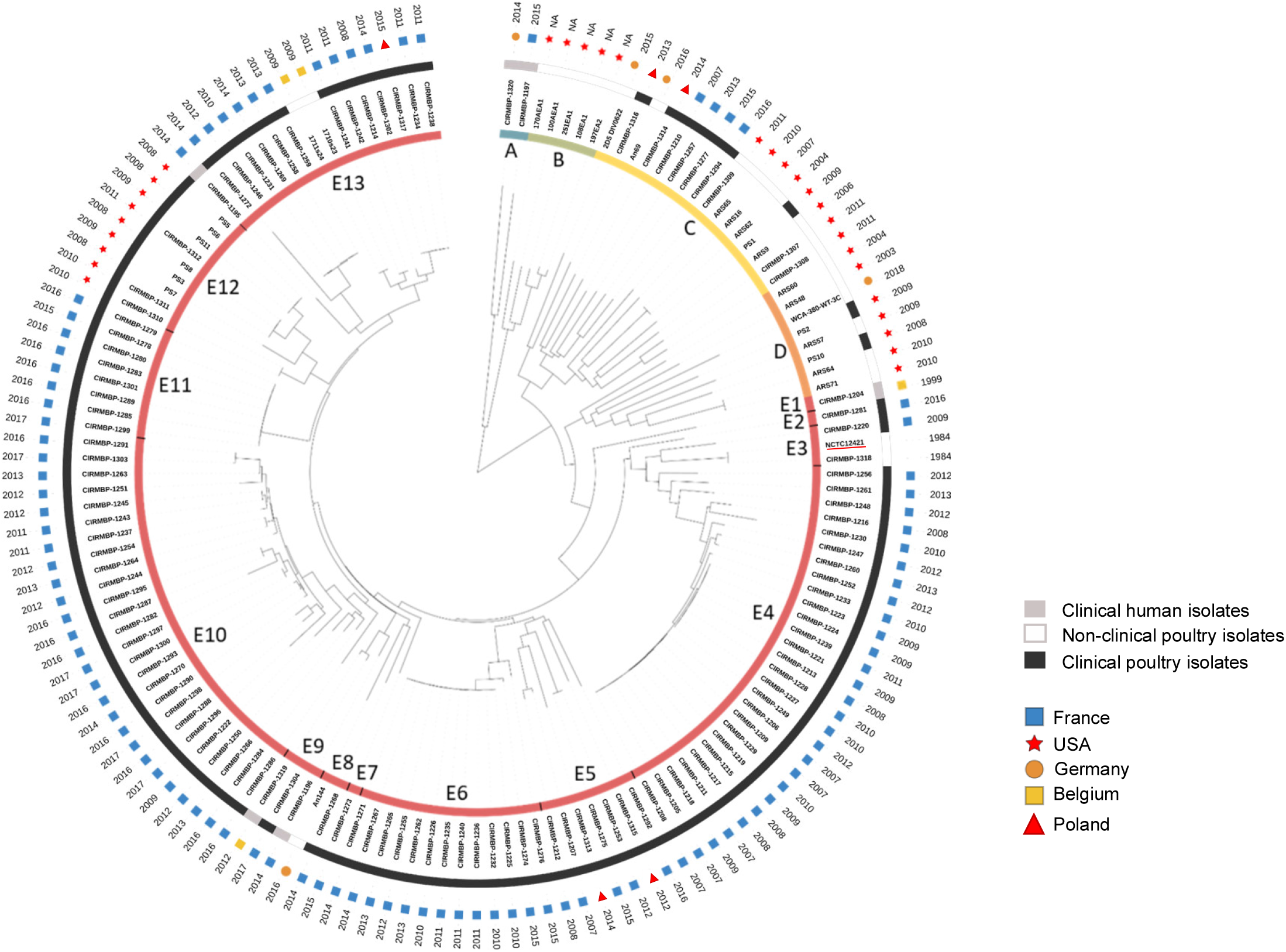
Phylogenetic tree and clinical status of 148 *E. cecorum* isolates. Neighbor-joining (BioNJ) tree built on pairwise distance between genomes. Internal circle: Clades A to E with subclades E1 to E13 (colored strips). First external circle: Clinical status of isolates (black: clinical poultry isolates, white: non-clinical poultry isolates, grey: clinical human isolates). Second external circle: geographic origin (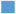: France, 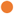: Germany, 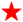: United States, 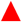: Poland, 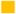: Belgium). Third external circle: year of isolation. The name underlined in red corresponds to reference genome NCTC 12421.

Altogether, these data revealed that clinical isolates of *E. cecorum* from poultry of different countries are grouped phylogenetically and mainly belong to clade E, confirming their clonality and suggesting either dissemination of well adapted clones or a convergent selection/adaptation to poultry genetics and breeding methods.

### Six genes, significantly associated with origin, differentiate avian clinical from non-clinical isolates in the 148 genomes

To further explore the genetic basis for the epidemic success of the *E. cecorum*, clade E isolates we searched for genes significantly associated with clade membership. The majority of unique genes (71.9%) are carried by isolates from clades A, B, C and D, indicating a higher genetic diversity in these clades. However, this trend may result from the enrichment of the collection in clade E isolates. Consistently, we identified 97 accessory genes significantly associated with clade E isolates (Table S3). The most abundant group of the 83 clade E-enriched genes after the unclassified hypothetical protein-encoding genes (n=17) are predicted carbohydrate metabolism and transport (n=13) genes and cell wall/membrane/envelop biogenesis genes (n=12). A high proportion of the clade E-enriched genes are clustered loci, including the capsule polysaccharide (CPS) biosynthesis locus (from CIRMBP1228_00568 to CIRMBP1228_00581 in strain CIRMBP-1228), a large gene cluster comprising biotin biosynthesis genes (CIRMBP1228_01807 to CIRMBP1228_01837), and one putative operon of carbohydrate metabolism (CIRMBP1228_02727 to CIRMBP1228_02733). Alignment of gene products of the representative capsule biosynthesis loci from the complete genomes identified two closely related loci represented by genomes of CIRMBP-1228 and CIRMBP-1212 (Fig. S3) that account for 66.7 and 14.5% of the clade E isolates, respectively. The remaining 14 genes significantly associated with origin of the isolates (clinical and non-clinical poultry isolates or clinical human isolates) are predominantly absent in clade E-isolates. They include genes encoding a pseudouridine synthase (CIRMBP1294_00547), a predicted nucleotide sugar dehydrogenase (CIRMBP1294_00386), a diacylglycerol kinase family lipid kinase (CIRMBP1294_00623), and two stress-related proteins (CIRMBP1294_00560 and CIRMBP1294_00758). In line with the large proportion of clade E clinical isolates in the collection, 30 clade E-enriched genes are also enriched in clinical poultry isolates compared to non-clinical poultry and clinical human isolates (Table S4A). They include genes of the CPS biosynthesis locus (CIRMBP1228_00573, CIRMBP1228_00574, CIRMBP1228_00575), 27 genes of the biotin gene cluster, and an H protein gene of the glycine cleavage system generally involved in protein lipoylation. A total of 65 genes are significantly associated with the origin of the isolates. In addition to those enriched in clade E isolates, other genes enriched in avian clinical isolates encode a phosphoenolpyruvate:carbohydrate phosphotransferase system (PTS), a transketolase (CIRMBP1228_00604 to CIRMBP1228_00607), as well as proteins of unknown function. Although no specific gene signature for avian clinical isolates was found, we identified 6 accessory genes (CIRMBP1228_00573, CIRMBP1228_00586, CIRMBP1228_00757, CIRMBP1228_01816, CIRMBP1228_02735, and CIRMBP1283_01819) that allow to identify 94 % of avian clinical isolates (Table S4B). With the exception of seven avian clinical isolates from clades C or D, all other genomes of clinical isolates have an average of 4 selected genes (range from two to six genes), while non-clinical isolates have one gene at most. We also found twelve genes that may be enriched in clinical human isolates. These include 8 genes predicted to be involved in import and utilization of ascorbate in anaerobic conditions (32, 33).

Despite the difficulty in identifying genes specific to the origin of the isolates, this work highlights six genes coding a glycosyltransferase (CIRMBP1228_00573), two PTS EIIC components (CIRMBP1320_01424, CIRMBP1228_02735,), and three hypothetical proteins (CIRMBP1228_00586, CIRMBP1228_01816, CIRMBP1228_00757) that can discriminate most of the isolates associated with poultry disease from those that are not.

### *Multiresistant clones of* E. cecorum *cluster in few clades*

Next, we searched the 148 sequenced genomes for the presence of acquired ARGs using ResFinder, BLAST and the PLSDB database. A total of eighteen ARGs were identified (Fig. 3). Tetracycline and macrolide-lincosamide-streptogramin B (MLS) resistance genes were detected in 95% and 75% of the isolates, respectively. In total, 70% of the genomes had at least one gene for resistance to these two families of antimicrobials, with *tet*(M) and *erm*(B) being the most prevalent. Resistance genes to other antimicrobial classes such as aminoglycosides (10%), bacitracin (30%), and vancomycin (0.6%) were also detected. Only two isolates (CIRMBP-1244 and CIRMBP-1314) did not contain any ARG searched. Overall, thirty-nine genomes carried genes conferring resistance to three antimicrobial families and seven genomes had at least four genes for resistance to different antimicrobial families (Fig. 3). The most common combinations of ARGs in multidrug resistant isolates cover the tetracycline and MLS families in combination with aminoglycosides in clades C and D or with bacitracin in clades C and D, and subclades E10, E11 and E12. Aminoglycoside resistance genes are carried by non-clinical genomes with an enrichment in U.S. strains. The near-systematic presence of ARGs in the genomes of *E. cecorum* isolates suggests that the species may have undergone strong selection for antibiotics, tetracyclines and MLS in particular, but that these two antibiotic resistances did not provide a selective advantage to clinical isolates.

**Fig. 3:**
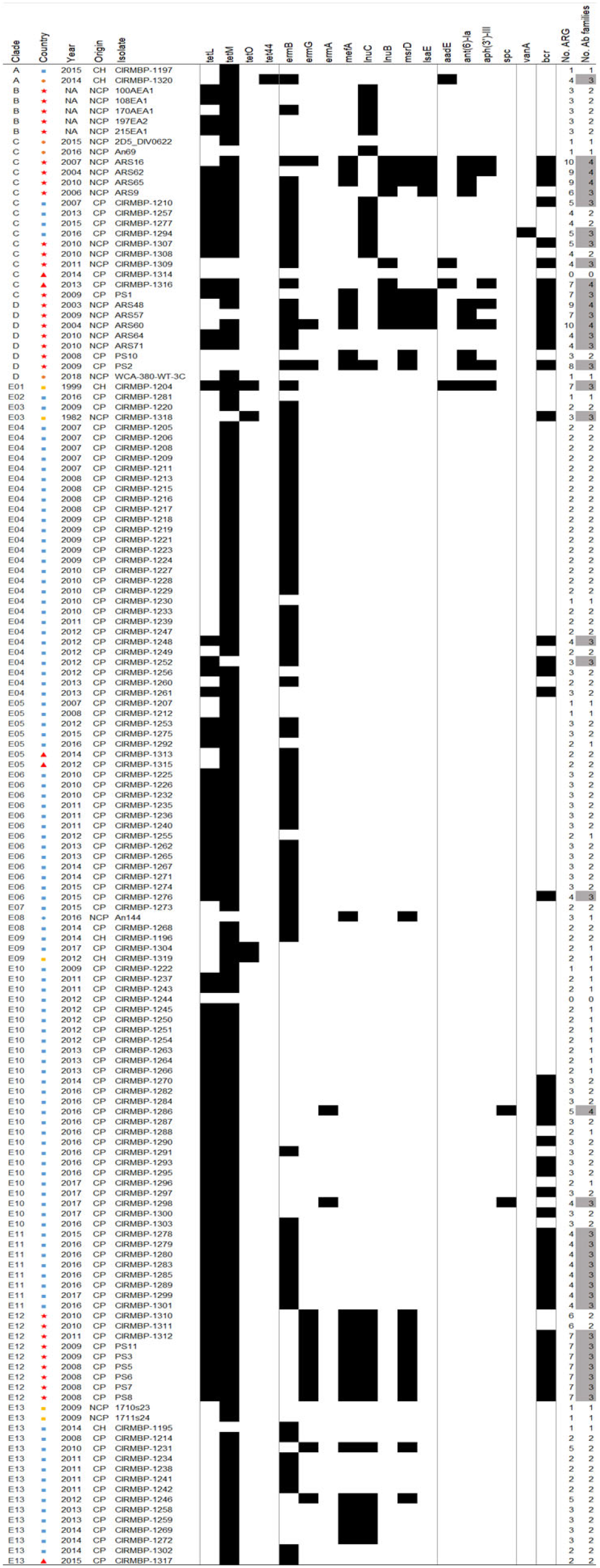
Distribution of antimicrobial resistance genes in sequenced genomes. Clade, geographic origin (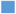: France, 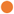: Germany, 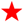: United States, 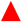: Poland, 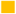: Belgium), isolation year and clinical origin of sample (CH: clinical human, NCP: non-clinical poultry, CP: clinical poultry isolates) are specified. Antibiotic families are represented by alternating grey blocks, from left to right: tetracycline, MLS (macrolide, lincosamide, and streptogramin), aminoglycoside, glycopeptide, and bacitracin. Black strips represent the presence of ARG and white ones the absence. Potential multiresistant isolates (>2 ARGs to different families) are highlighted in grey.

Several *E. cecorum* genome regions carrying ARGs conserved in Gram-positive bacteria were identified in the PLSDB database, but none was encoded by a previously identified plasmid. Their analysis highlighted different ARGs clusters. They comprised *erm*(B), *vat*(D), and *msr*(D) genes, *tet*(L), *tet*(M), and *bcr* genes or *tet*(L), *tet*(M), *ant(6*), and *bcr* genes. Further characterization of the vancomycin resistance operon carried by strain CIRMBP-1294 revealed that the *vanA* operon was integrated into the chromosome 12 kbp away from the genes that confer resistance to narasin and to erythromycin in between (Fig. S4). This chromosomal region was closely related to that of the *E. faecium* plasmid pVEF3 (34) identified in broiler isolates from Sweden (35) and was flanked by two transposase genes of the IS*3* family. However, no conjugative or mobilization element was detected close to this ARG cluster. Although about twenty plasmid-related gene families were identified in the pan-genome, a single putative 4.4 kbp plasmid was detected in the complete genome of strain CIRMBP-1292 and 6 other isolates (CIRMBP-1233, CIRMBP-1259, CIRMBP-1286, CIRMBP-1289, and CIRMBP-1304), but no ARG was detected on this small plasmid. Together, these results suggest that plasmids are rare in this bacterial collection and ARGs are encoded on mobile genomic islands.

### E. cecorum *mobilome is a major contributor to inter- and intra-clade diversity*

In Gram positive bacteria, ARGs and related transposons are frequently integrated in complex MGEs forming Integrative Conjugative Elements (ICEs) or Integrative Mobilizable Elements (IMEs) that may be difficult to identify in draft genomes (36). To evaluate the contribution of these elements to the spread of ARGs and to *E. cecorum* genome diversity, we predicted ICEs and IMEs in the eleven complete genomes and the two large contigs of genome CIRMBP-1320 using ICEscreen (J. Lao, T. Lacroix, G. Guédon,C. Coluzzi, N. Leblond-Bourget, and H. Chiapello, submitted for publication). ICEscreen ICE and IME detection relies on the presence of Signature Protein CDSs (integrase, coupling protein, relaxase, and virB4) grouped on the genomes and previously shown to be a valuable clue to the presence of an integrative element (37, 38). ICEs were defined by the superfamily and family of their signature proteins (37). Each *E. cecorum* genome contained at least one ICE (Table S5A and Fig. 4). In total, thirty-three mobile genetic elements including five types of ICEs (belonging to the *Tn916, Tn5252*, and *TnGBS1* superfamilies), two IMEs, three *Tn917*, and one partial conjugative element were identified. Their size ranged from ~5 kbp for Tn*917* to ~103 kbp for the *TnGBS1* ICE of genome CIRMBP-1302. The Tn916 ICE of genomes CIRMBP-1228 and CIRMBP-1281, integrated upstream of the 30S ribosomal protein S6 encoding gene, included another ICE of undescribed family encoding a DDE transposase, a MobC-like relaxase, a VirD4-like coupling protein, and a type IV secretion system (T4SS) protein VirB4. We also investigated the presence of other genomic islands (GIs) in the complete genomes (Fig. 4, Table S5B). A total of 42 GIs were identified, with a size ranging from ~8 kbp to ~122 kbp for the complex GI that comprises an ICE (Tn*916*, ICE*Bs1*, ICE*St3*) in genome CIRMBP-1228. Each genome harbored from 3 ICE-related elements or genomic elements (strain CIRMBP-1212 of subclade E5) up to 12 (strain CIRMBP-1228 of subclade E4). They were mainly integrated in intergenic regions or the 3’-end of genes with no effect on the encoded sequences. As expected, all Tn*917* and ICEs of the Tn*916* family encoded *erm(B*) and *tet*(M), respectively. Other ARGs such as *tet*(L), *tet*(O), *ant(6*), and the *bcr* operon were carried on Tn*916*-related ICEs or on GIs. Besides genes involved in GI transfer, a few had predicted functions related to biotin biosynthesis, restriction modification enzymes, cadmium/arsenate resistance, toxin-antitoxin systems, redox enzymes, type 2 secretion system, and flagella and chemotaxis. Analysis of the distribution of the ICE-related elements and the GIs in the other 136 *E. cecorum* genomes (Table S5B) revealed three highly dispersed elements: the Tn*916*-related ICE of strain CIRMBP-1292 in 113 genomes, the GI inserted near the *rpsB* gene and encoding the biotin biosynthesis genes found in 90 genomes, and the transposon *Tn917* in 65 genomes (Table S5B). The GI inserted between *dnaX* and *sufB* of strains CIRMBP-1287 and CIRMBP-1274 was highly prevalent in subclades E6 and E10, respectively. The same is true for the cognate element of CIRMBP-1283 and CIRMBP-1302 that is detected in all isolates of subclade E11, to which CIRMBP-1283 belongs. Other elements are enriched in specific subclades, such as the GI near the 23S rRNA methyltransferase *rlmD* gene and ICEs of the *TnGBS1* and *Tn916* families of CIRMBP-1274 enriched in subclade E6, the ICE of Tn*916* family of CIRMBP-1212 enriched in subclade E13, the GI inserted in the 3’-end of CIRMBP1228_01030, and the ICE of the *TnGBS2* family of CIRMBP-1228 close to the gene CIRMBP1228_01622 enriched in E4. According to their gene content, fifteen elements appear to be more strain-specific in strains CIRMBP-1320 (n=4), CIRMBP-1281 (n=4), CIRMBP-1292 (n=3) and NCTC12421 (n=4). Although the genomes used to identify ICEs and GIs are not representative of all clades, homologs were found in almost all clades except clade B, which gathers only five non-clinical poultry isolates from the United States. Yet However, these genomic elements were less conserved in isolates from the United States, suggesting they were acquired independently. We conclude that the majority of the identified ICEs and GIs are enriched in some clades, while only a few are shared across clades.

**Fig. 4:**
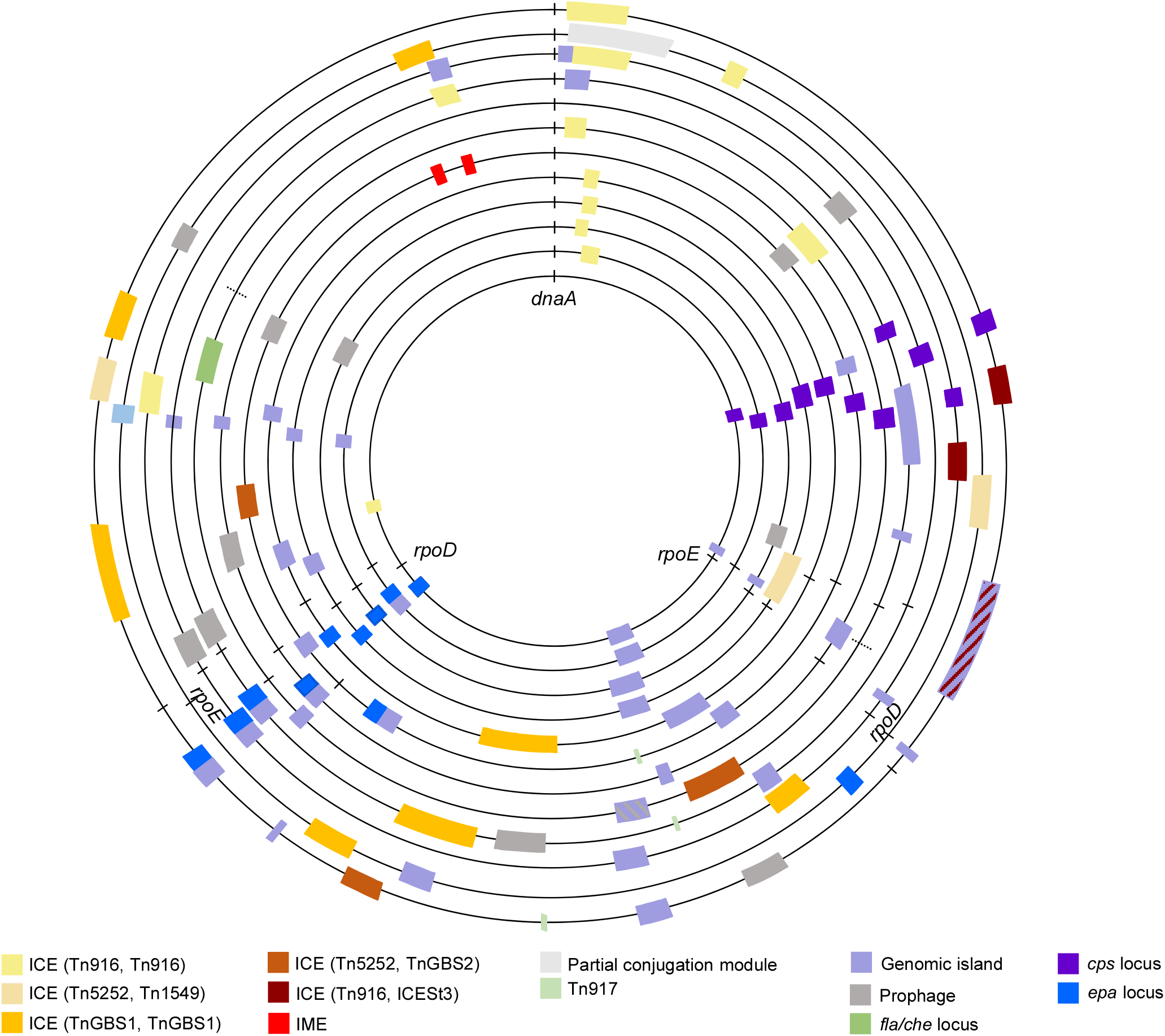
Mobile genetic elements in complete *E. cecorum* genomes. From inner to outer circle: CIRMBP-1212, CIRMBP-1283, CIRMBP-1292, CIRMBP-1246, CIRMBP-1302, NCTC12421, CIRMBP-1287, CIRMBP-1320, CIRMBP-1274, CIRMBP-1281, CIRMBP-1261, CIRMBP-1228. Dotted lines on genome CIRMBP-1320 correspond to predicted junctions. Each colored rectangle indicates the integration of an element according to the color legend. Hatching indicates a complex genomic island. *dnaA*, *rpoE*, *rpoD* genes and *epa* and *cps* loci are indicated.

As prophages are easier to detect than GIs and mobile genetic elements, prophages were searched for in all *E. cecorum* genomes sequenced, including NCTC12421. A total of 103 complete prophages and 30 incomplete prophages were identified in 78 (66.1 %) *E. cecorum* genomes (Table S6). Prophage size ranged between 31 and 59 kbp in length. The prophages were related to 19 different types that varied across the phylogenetic groups. The subclade E4 was enriched with prophages related to *Faecalibacterium* phage oengus (39) that was the most commonly found (n=31), followed by prophages related to the *Siphoviridae* temperate phage EFC-1 of *Enterococcus faecalis* (40) (n=16). Prophages related to the *Myoviridae* temperate bacteriophage EJ-1 of *Streptococcus pneumoniae* (41) (n=13) were mostly found in genomes of subclade E10 while those related to the *Siphoviridae* temperate phage of *Lactococcus lactis* 50101 (n=12) were most prevalent in subclade E6. Conversely, clade C had the most diverse prophages and subclades E10 and E12 contained the least number of prophages. Prophages related to *Siphoviridae* temperate phage 5093 of *Streptococcus thermophilus* (42) (n=9) were identified in several phylogroups. A total of 13 different integration sites for temperate bacteriophages were identified, mainly in intergenic regions and at the 3’ end of genes without changing the reading frame (Table S6). However, the prophage related to PHAGE_Strept_EJ_1_NC_005294 integrated in the PreQ1 riboswitch is likely to change the expression of the downstream nucleoside hydrolase gene. Like ICEs and GIs, prophages are less prevalent in the ~700 kbp surrounding the origin of replication of the chromosome (Fig. 4), indicating that mobile genetic elements are not randomly distributed in the *E. cecorum* genome.

### Biofilm robustness, adhesion to type II collagen, and growth in chicken serum are not associated with gene content

To get phenotypic insight into the 118 isolates, we independently examined phenotypes relevant to *E. cecorum* pathogenesis: biofilm robustness, adhesion to type II collagen, and growth in chicken serum (see methods in the supplemental material). Hierarchical clustering of these phenotypes (Fig. 5) revealed 17, 11 and 9 groups of strains for biofilm robustness, type II collagen adhesion, and growth in chicken serum, respectively. The lack of concordance between these phenotypic groups and clades strongly suggests that none of the phenotypes are discriminating between clades. In an attempt to rank isolates, the most robust biofilm-forming isolates were CIRMBP-1302 (subclade E13), CIRMBP-1277 (clade C) and CIRMBP-1228 (subclade E4) and the most fragile biofilm-forming isolates were CIRMBP-1204 (subclade E1), CIRMBP-1205 and CIRMBP-1211 (subclade E4), and CIRMBP-1225 (subclade E6). The most collagen-adherent isolates were CIRMBP-1274 (subclade E6) and CIRMBP-1277 (clade C). All of the *E. cecorum* isolates had a serum growth index below the index of the *Escherichia coli* control strain sensitive to the bactericidal effect (190.34 +/- 123.88) of the serum and in the range of the non-sensitive *E. coli* control strain (3.172 +/- 4.2). The *E. cecorum* isolates with the lowest indexes were strains CIRMBP-1318 (0.229 +/- 0.21) and CIRMBP-1206 (0.467 +/- 0.001) belonging to subclades E3 and E4, respectively. The *E. cecorum* strains with the highest indexes were strains CIRMBP-1312 (3.85 +/- 0.44) of subclade E12, CIRMBP-1197 (4.21 +/- 2.2) of clade A, and CIRMBP-1298 (5.71 +/- 2.9) of subclade E10. Taken together these results indicate that the growth of *E. cecorum* is not affected by the presence of chicken serum. Beside strain CIRMBP-1277, which formed strong biofilms and had a high capacity to adhere to collagen, no relationship between the expression of the three phenotypes and adherence to collagen was observed. Moreover, no accessory genes were associated with the phenotypic groups or with binary transformed values, suggesting functional redundancy between genes and/or differential gene expression between isolates.

**Fig. 5:**
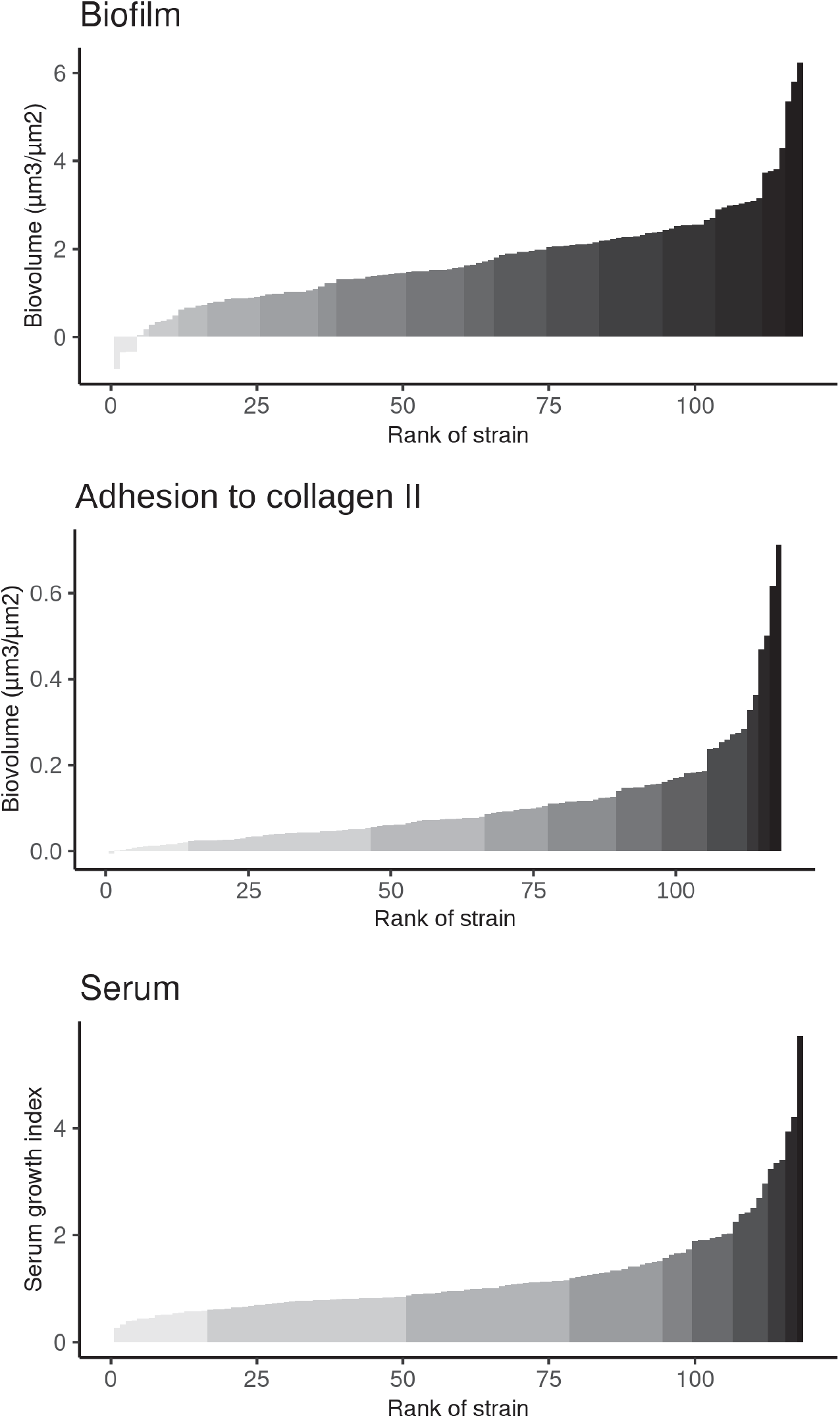
Distribution of phenotypic expression for biofilm robustness, adhesion to type II collagen, and growth in chicken serum for 118 *E. cecorum* isolates. Values for biofilm and adhesion to collagen represent estimated marginal mean biovolumes. Values for growth in serum represent estimated marginal mean of serum growth index calculated at 6 h (see supplemental material). Each bar corresponds to an isolate. Strains with similar phenotypic expression were grouped into clusters in grey color scale.

### *Clinical isolates of* E. cecorum *show a broad spectrum of virulence in chicken embryos*

To evaluate the pathogenic potential of *E. cecorum* strains isolated from diseased broilers in French farms, we used the chicken embryo lethality assay (CELA) reported by Borst et al. using strain CIRMBP-1309 (original name CE3) and strain CIRMBP-1311 (original name SA2) as biological controls for a non-pathogenic and pathogenic poultry isolate, respectively (21). We tested the 11 *E. cecorum* strains with the most complete genomes (see methods in the supplemental material). No embryo died in the control group. The two *E. cecorum* control strains behaved as expected: 53 % of the embryos were still alive 6 days after inoculation with the commensal strain CIRMBP-1309, whereas 100% of embryos died after 2 days when inoculated with strain CIRMBP-1311 isolated from spondylitis lesions (Fig. 6). Strain CIRMBP-1294, isolated from infected vertebrae and strain CIRMBP-1320, isolated from human infection induced less than 27% embryonic lethality, indicative of a low virulence in CELA. In contrast, strains CIRMBP-1228, CIRMBP-1274, CIRMBP-1292, CIRMBP-1302, and CIRMBP-1304, mainly isolated from infected vertebrae, showed the same virulence as strain CIRMBP-1311 with an overall average lethality of 85% at day 2 post-infection. They thus could be considered as virulent isolates. The lethality of strains CIRMBP-1212, CIRMBP-1281, CIRMBP-1283, and CIRMBP-1287 was intermediate and higher than observed for the least virulent strains CIRMBP-1294 (p-values between 0.026 and 0.064) and CIRMBP-1320 (p-values between 0.026 and 0.073). Close analysis of the gene content in the 10 strains exhibiting virulence and in the three strains considered as non-virulent in CELA identified 33 genes, of which 18 were already found enriched in clade E isolates and 6 in avian clinical isolates. Predicted function for these genes indicates a bias in favor of carbohydrate transport and metabolism (Table S7).

**Fig. 6.**
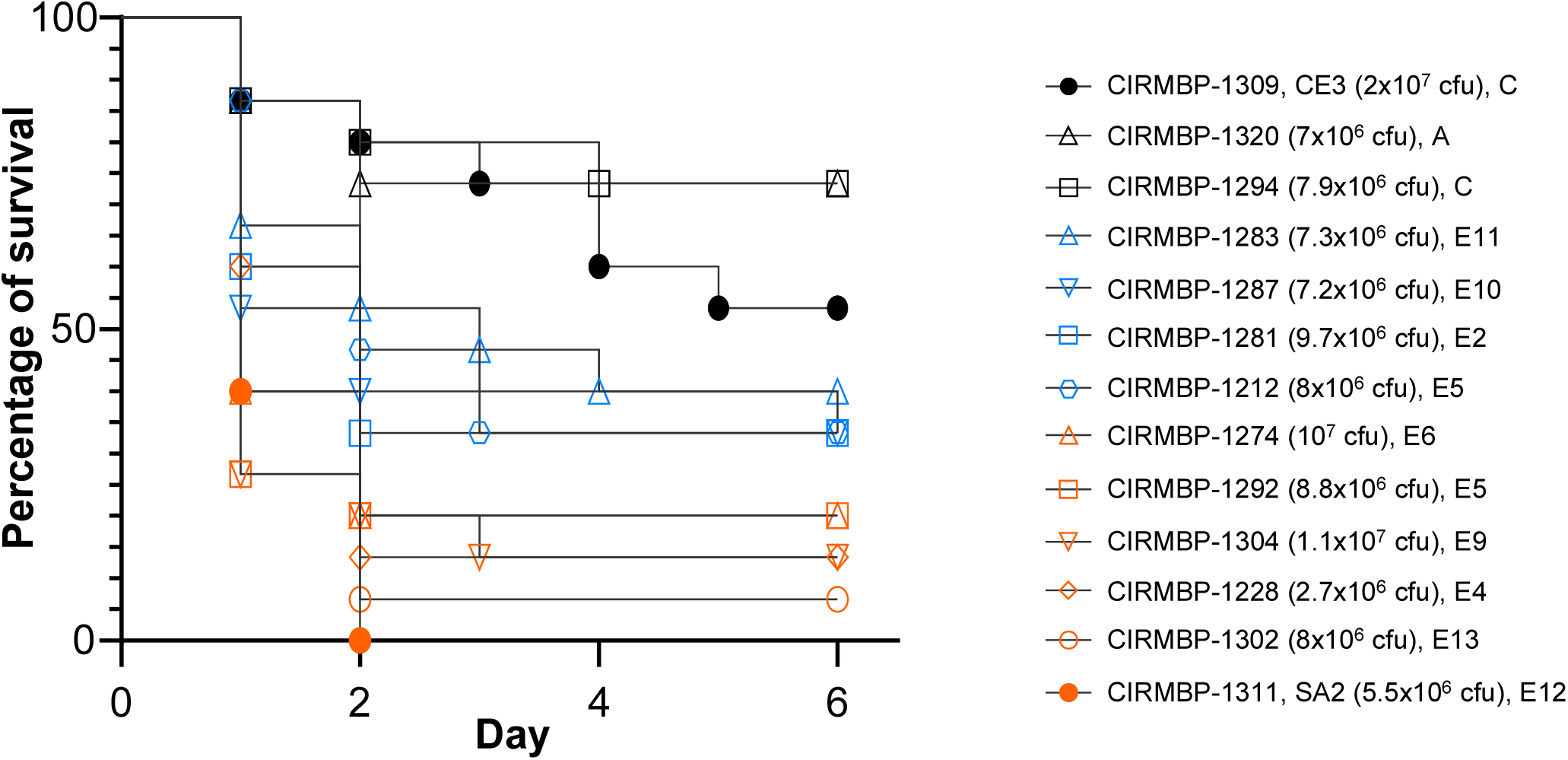
Comparison of virulence of selected *E. cecorum* isolates in a chicken embryo model of infection. Kaplan–Meier survival plot of chicken embryos (n=15) infected with 13 different isolates of *E. cecorum*; strain name, inoculum size, and phylogenetic group are indicated. The log-rank (Mantel-Cox) test indicated significant differences between the positive control CIRMBP-1311 (SA2) and strains CIRMBP-1309 (CE3, p-value <0.0001), CIRMBP-1212 (p-value= 0.0007), CIRMBP-1281 (p-value = 0.0364), CIRMBP-1283 (p-value = 0.0035), CIRMBP-1287 (p-value = 0.0339), CIRMBP-1294 (p-value <0.0001), and CIRMBP-1320 (p-value <0.0001). Virulence of strains CIRMBP-1292, CIRMBP-1304, CIRMBP-1228, CIRMBP-1274, and CIRMBP-1302 was not significantly different from CIRMBP-1311. One representative experiment of two is shown.

In line with data obtained for biofilm robustness, binding to collagen, and growth in chicken serum, the clear gradient of virulence observed in the CELA for clinical isolates, supports the hypothesis of the multifactorial nature of *E. cecorum* pathogenesis.

## DISCUSSION

*E. cecorum* has emerged as an opportunistic pathogen in poultry worldwide. The present study shows the clonality of a large collection of clinical *E. cecorum* isolates collected between 2007 and 2017 in the leading French commercial broiler producing area. It identifies main phylogenetic clades and subclades and provides a first insight into the intercontinental clonality of clinical isolates of *E. cecorum* from poultry. Six genes significantly associated with the origin of the isolates allow to discriminate 94% of the avian clinical isolates of the collection from the non-clinical ones. Based on new complete genomes, we also provide insight into the diversity of the mobile genetic elements of *E. cecorum* that carry ARGs.

One way of assessing the diversity of a species is to analyze its core- and accessory-genomes (43–45). Although it should be considered as rough estimates due to the use of draft genomes, the current *E. cecorum* core-genome represents only 14.1% (1,207/8,523 CDSs) of the pan-genome and ~50% of the average *E. cecorum* genome. While the size of the core-genome is within the range of the one reported by Sharma et al., the *E. cecorum* pan-genome is now 35% larger than in earlier estimates due to the addition of 130 genomes to the 18 genomes previously used (28). The increase in genome size corresponds to an average of only 20 genes per genome; however, the low gene discovery rate is probably due to the high proportion (93%) of clinical poultry isolates in clade E. Indeed, clade E isolates (n=117) accounted for 22% (1,873/8,523) of accessory genes while more distantly related isolates of clades A (n=2), B (n=5), C (n=16) and D (n=8) accounted for 34.0% (2,897/8,523) of accessory genes. The proportion of core genes and accessory genes strongly correlates with the lifestyle of the bacterium. A small core-genome compared to a large pan-genome reflects the diversity of hosts and lifestyles, as observed in *E. coli sensu stricto* and *Salmonella enterica*, two ubiquitous species with commensal or pathogenic lifestyles, whose core-genome accounts for 0.39% and 1.9% of the pan genome, respectively (46, 47). Conversely, a high proportion of core genes reflects more restricted lifestyles, such as those of *Bacillus anthracis* (65%), an obligate pathogen and *Staphylococcus aureus* (36%) and *Streptococcus pyogenes* (37%), two human-restricted pathogens (44). Broader sampling from different hosts and countries is needed to further evaluate the diversity of the *E. cecorum* species.

Genome reduction associated with pathogenicity is observed in many bacteria, including *Streptococcus suis* and *Streptococcus agalactiae* whose genome size is reduced in virulent host-adapted isolates (48–51). Although longer *E. cecorum* genomes belonged to clinical poultry isolates, the average genome size was similar for all poultry isolates regardless of their clinical status (2.40±0.14 and 2.36±0.11 Mbp for clinical poultry isolates and non-clinical poultry isolates, respectively). This observation differs slightly from two previous studies on isolates from the United States, where clinical isolates had shorter genomes than non-clinical isolates (19, 28). The latter is most likely due to the small number of genomes examined and the sampling of clinical isolates, which according to our findings belong all to the same phylogenetic subclade E12. In the current dataset, the smallest genomes correspond to isolates from phylogenetic subclade E11 and the largest genomes to isolates from subclade E4. Interestingly, all but one E4 genomes (n=27) have no CRISPR-*cas* systems, correlating with abundant ICEs, GIs and prophages.

Previous molecular studies, using mainly PFGE, have converged towards the genetic homogeneity of *E. cecorum* clinical isolates from the same country compared to non-clinical isolates (19, 22–28). Despite more than 6,500 cases of *E. cecorum* infections in poultry reported in France since 2007 (52), the genetic diversity of *E. cecorum* in France and genetic relatedness with isolates from other countries have never been studied. Phylogenetic analysis of 100 isolates spanning the period from 2007 to 2017 showed that a single clade (E) was responsible for almost all (96%) cases in farms between 5 and 90 km apart in Brittany. The phylogenetic congruence of clinical isolates from the United States is consistent with two hypotheses: i) isolates were issued from a transcontinental dissemination or ii) isolates from different regions suffered comparable selective pressure. The use of a limited number of commercial genetic lines of broiler chickens may have contributed to the selection of *E. cecorum* clones with pathogenic potential and allowed transcontinental dissemination. While the hypothesis of a transcontinental spread of clade E isolates through the meat trade is unlikely, trade in live animals (53), surface-contaminated eggs, or transport by wild birds may well have contributed to this spread. The second hypothesis, although less likely, is a parallel convergent selection due to breeding conditions. The isolates in this study may not represent the full diversity of the French clinical population of *E. cecorum;* however, their temporal distribution reflects the spread and persistence of a clade particularly adapted to broilers with the emergence over time of some more successful subclades. This is illustrated by the dominance of subclade E4 (82%) in France and subclade E12 (75%) in the United States between 2008 and 2010 in the current dataset. Subsequently, the dominant subclades were E6, E13, E11 and E10, the latter two were dominant on French farms in 2016. The temporary circulation of specific subclades may be due to natural evolution or adaptive changes in response to modifications of breeding practices (like novel biocides and cleaning procedures, different feed origin, composition, or additives), but the reasons remain to be determined. Though less frequent, clinical isolates were also found in clades C and D, which contain non-clinical isolates and display higher diversity. Noticeably, isolates of these two clades have multiple ARGs, which may confer selective advantage and thus contribute to the pathogenic potential in specific conditions that remain to be determined. Additional genomes of non-clinical isolates, but also clinical isolates from diverse countries and other poultry species and husbandry systems are required to obtain a comprehensive view of the *E. cecorum* population structure and determine whether *E. cecorum* clade E isolates are broiler specific.

*E. cecorum* has been occasionally involved in human infections (54–56). Four of the six clinical human isolates of this study were clade E isolates, supporting a poultry origin. However, this does not lead to the conclusion that the contamination was food-borne. The two other clinical human isolates belong to clade A and are phylogenetically close. Both have large clusters of highly specific genes, including an iron transporter and a ~60 kbp motility locus characterized by flagella and chemotaxis genes, which may have been acquired by horizontal gene transfer from other species of enterococci, such as *E. casseliflavus*, *E. gallinarum*, or *E. columbae* encountered in birds, but also in other animals including humans, insects, and aquatic hosts for the first two (57).

The dominance of *E. cecorum* clinical isolates from clade E strongly supports the hypothesis that clade E isolates have acquired properties increasing their fitness and/or infectivity. At the core-genome level, all clade E isolates share common SNPs that may confer a selective advantage in the host. Of the two non-neutral mutations one is in the phage shock protein gene *pspC* that encodes an ortholog of the transmembrane protein LiaY, probably involved in resistance to cationic antibiotics and antimicrobial peptides (58). Codon changes leading to synonymous SNPs may also modify the translation efficiency as such to foster cell fitness (59). In addition to the clade E-specific mutations in the core genes, 83 accessory genes were found enriched in the clade E isolates of which thirteen genes of the capsule operon. The capsule is an important virulence factor to evade host immunity, including for enterococci (see (60) for a review on the subject). We identified two closely related capsule loci in 81.2% of the clade E isolates. The strong association between these capsule loci and the clinical isolates suggests a role in virulence, probably by promoting immune evasion (61). However, virulence, especially for opportunistic pathogens, is a multifactorial process involving multiple bacterial traits such as metabolic functions and stress resistance (62, 63). Other clade E enriched genes may confer *E. cecorum* alternative metabolic capacities to survive and/or multiply in the host. The predicted galactitol phosphotransferase system (CIRMBP1228_02729 to CIRMBP1228_02731) and galactonate catabolism enzymes (CIRMBP1228_02728, and CIRMBP1228_02732), as well as the biotin biosynthesis genes may give *E. cecorum* an advantage in competing with commensal species in nutrient-limited environments such as the gastro-intestinal tract, considered as a portal of entry during the first week of life of the host (10). Of note, the mannitol-1-phosphate 5-dehydrogenase (*mtlD*) gene proposed to be specific of non-clinical isolates (19) was not discriminant between non-clinical and clinical poultry isolates of our collection because it is present in only seven isolates from different origins. Conversely, we selected 6 genes, which taken together allow to discriminate more than 90% of the clinical isolates. The combined detection of these candidate genes on a larger collection of clinical and non-clinical isolates is necessary in order to evaluate their use for the detection of early carriage of potentially pathogenic isolates, particularly during the first week of life. In contrast, while ascorbate catabolism genes are dispensable in 70% of clinical avian isolates, they may confer a competitive advantage to avian non-clinical isolates and human clinical isolates.

MGEs including prophages are major contributors to the evolution of the gene repertoire. We identified 75 MGEs corresponding to predicted ICEs or related elements and GIs in the complete sequenced genomes and a total of 105 complete prophages in the 118 sequenced genomes. Three GIs have a composite structure with phage genes and more than one integrase gene, likely resulting from independent integration of different MGEs (37, 64). *E. cecorum* MGEs are integrated in the 3’ end or in intergenic regions of genes encoding a ribosomal protein (*rpsB, rpsI, rpsF*, or *rpmE*), in a few tRNA genes or riboswitches (*tRNA-Thr, tmRNA, preQ1*), but also in various other intergenic regions. This is consistent with the different site-specificity of the prevalent integrases of the tyrosine and DDE recombinase families (36–38), although there is relatively little integration in the tRNA genes that are frequently targeted by tyrosine integrases in streptococci (65). The apparently non-random distribution of the *E. cecorum* MGEs integrated relatively far away from the chromosomal origin of replication may relate to the eviction of highly expressed genes located near the origin of replication (66, 67). Among the prophages we identified, prophages homologous to PHAGE_Strept_5093, PHAGE_Entero_EFC_1, and PHAGE_Bacill_phBC6A52 were also detected in clinical and non-clinical poultry isolates from the United States (28). In contrast, and with the exception of Tn*917*, the ICEs and GIs identified in the French isolates are not well conserved in the U.S. isolates, indicating that they were acquired separately and contribute to local adaptation. In addition to the type IV secretion system involved in the formation of the DNA translocation channel, ICEs encode cell surface adhesins for attachment to the target cell. These include LPxTG cell wall-anchored adhesins such as *S. agalactiae* antigen I/II family adhesins also referred to as group B *Streptococcus* surface proteins (Bsp) (68, 69). These structural proteins were also shown to promote biofilm formation, interaction with host cells, and virulence (70, 71). We identified Bsp-like proteins inTn*GBS1* and Tn*GBS2*-related ICEs, proteins containing Cna B domains initially found in microbial surface components recognizing adhesive matrix molecules (MSCRAMMs), and VaFE repeat-containing surface-anchored proteins in diverse *E. cecorum* ICEs and GIs. Yet none of these associated with a specific trait or origin. However, the variability of these adhesins may hinder any association, as they may also compensate each other. ICEs and GIs may carry diverse genes, known as cargo genes, that are not involved in gene transfer, but may confer a selective advantage to the host strain. We have identified several *E. cecorum* cargo genes encoding toxin-antitoxin modules that function as MGE addiction systems but are also involved in the control of bacterial growth (72). Other cargo genes encode restriction-modification systems that protect the cell against horizontal gene transfer or are genes involved in protection against oxidative stress, or in resistance to cadmium or arsenate, which could confer a better fitness contributing to ecological adaptation. An accessory SecA2-SecY2 operon was also identified in strain CIRMBP-1228. Such systems are dedicated to the export of glycosylated serine-rich repeat proteins (SRRPs) that participate in adhesion to host cells and/or in biofilm formation (73, 74). Functional analysis on a few selected strains is required to evaluate whether and how these accessory genes contribute to adaptation to environmental challenges.

ARGs are other clinically important cargo genes spread by ICEs and GIs (75). The ARGs identified in *E. cecorum* confer resistance to tetracycline, macrolides, bacitracin, aminoglycosides, and much more rarely to glycopeptides. The most prevalent are *tet*(M), *tet*(L), and *erm*(B) genes. This is in line with the high prevalence of resistance to tetracycline and erythromycin in *E. cecorum* found in various studies, as reviewed by Jung et al. (12) and with the use of tetracyclines and macrolides despite substantial efforts to reduce their use in veterinary medicine. As anticipated from the literature, *tet*(M) is carried on ICEs of the Tn*916* family and *erm*(B) on Tn*917* (76). Other macrolide resistance genes, such as *mef*(A), *msr*(D), or *lnu*(B) and the aminoglycoside resistance genes, such as *ant(6)-Ia* and *aph(3 ‘)-III* are prevalent in the USA U.S. isolates (28). *mef*(A), *msr*(D), *vat*, and *erm*(C) genes are located on the same GI in CIRMBP-1246 (CIRMBP1246_01012-CIRMBP1246_01050), *lnu*(C) is adjacent to the IS*1595* family transposase *ISSag10* and the two adjacent genes *lnu*(B) and *lsa*(E) are next to the IS*1595* family transposase *ISCpe8*, previously described in an avian *Clostridium perfringens* strain carrying the lincomycin resistance gene *lnu*(P) on the plasmidic transposable tIS*Cpe8* (77). Another prevalent ARG is the bacitracin resistance operon *bcr* in isolates of clades C, D, and subclades E10, E11, and E12. This operon is frequently associated with *tet*(M) and *tet*(L) genes on ICEs of the *Tn916* family. The highly conserved nucleotide sequence of the *bcr* operon, including the flanking IS*Enfa1* and its location on Tn*916*-like elements or GIs, is consistent with avian inter-species transmission involving *E. faecalis, E. faecium* and *C. perfringens* (78, 79). Note that the carriage of *tet*(M) and *tet*(L) on the same Tn916-like element is uncommon. It was first described in *Streptococcus gallolyticus* and proposed to benefit the host bacterium under stressful conditions (80, 81). A single clinical poultry isolate from France has the *vanA* operon, a gene already described in an *E. cecorum* strain from retail poultry in Japan (82). Overall, 26% (n=39) of isolates encode multiple ARGs (4 to 10) conferring resistance to at least three antimicrobial families, and are prevalent in clinical and non-clinical isolates of clades C, D, and in subclades E11 and E12. The very few strains without predicted ARG and the differential ARG profiles between French and U.S. isolates probably reflect a strong antibiotic selective pressure that differs between the two countries. Indeed, in-feed bacitracin and in-feed macrolides are still used in poultry farming in the United States (83). The successive European bans of antibiotics (avoparcin in 1997, bacitracin, spiramycin, tylosin, and virginiamycin in 1999, avilamycin and flavophospholipol in 2006) and the French national EcoAntibio plans (84, 85) launched in 2012 and 2017 to fight antimicrobial resistance in animal health and promote the responsible use of antibiotics might have contributed to contain the spread of ARGs and reduce MLS resistance genes as observed in the recent isolates of subclade E10. In fact, this is in line with the decreasing trend of macrolide resistance of *E. cecorum* strains isolated in French poultry according to the French surveillance network for antimicrobial resistance in bacteria from diseased animals (RESAPATH on line, https://shiny-public.anses.fr/resapath2/). However, there has been a marked increase in bacitracin resistance genes in French isolates since 2015, even though bacitracin is not used in avian veterinary medicine in France (86). Bacitracin is produced by *Bacillus licheniformis* and *Bacillus subtilis* strains. With the need for alternatives to antibiotics in livestock, bacilli strains are used as probiotics or applied together with lactic acid bacteria as protective biofilm against pathogens (87–89). The increase of *E. cecorum* isolates carrying the *bcr* operon points to the need for examining whether *Bacillus* strains applied in farms produce bacitracin or related antimicrobial compounds that could contribute to the dissemination of the *bcr* operon. Reassuringly, relatively few aminoglycoside and vancomycin resistance genes that target gentamicin and glycopeptides, two critically important antimicrobials in human medicine, are found in French isolates, as elsewhere in Europe (12).

Overall, the results of this study shed light on the population of *E. cecorum* clinical isolates in France and reveal a genetic linkage with *E. cecorum* clinical isolates from elsewhere. We have shown that, based on the available data, the majority of clinical poultry isolates are phylogenetically distinct from non-clinical poultry isolates and form a main clade responsible for the outbreaks of *E. cecorum* in France and probably in the United States and Europe. ICEs and GIs are the main carriers for antimicrobial resistance. The E clade of *E. cecorum* appears to have adapted to the conditions under which poultry is reared, highlighting its importance as an emerging threat to the poultry industry worldwide. This information can be used to design and guide preventive strategies to reduce the impact of *E. cecorum* clade E isolates.

## MATERIAL AND METHODS

### Bacterial strains

A total of 118 strains were collected from various laboratories and deposited at the International Center for Microbial Resources-Bacterial Pathogens (CIRM-BP, https://www6.inrae.fr/cirm_eng/BRC-collection-and-catalogue/CIRM-BP) (Table S1). The majority of them (n=100) were isolated between 2007 and 2017 from diseased birds in 16 broiler farms located in Brittany, France and were provided by Labofarm (Loudéac). Other poultry strains were isolated in Poland (n=5), Belgium (n=1), and the United States (n=6). Six strains from human infections were isolated in France (n=3), Belgium (n=2), and Germany (n=1). The source and the original name of the strains from abroad is indicated in Table S1. Additional details are available in the supplemental material.

### Genome sequencing and analysis

All genomic DNA was subjected to random shotgun library preparation using the TruSeq DNA PCR-Free kit (Illumina). Ready-to-load libraries were sequenced on Illumina Miseq or HiSeq 3000 platforms (Illumina) at GeT-PlaGe (Toulouse, France) and HiSeq 2500 platform at Eurofins Genomics (Germany) using 150 bp paired-end chemistry. DNA of fourteen isolates was also sequenced using Oxford Nanopore Technology platforms. Preparation of libraries and sequencing were performed at the GeT-PlaGe core facility (INRAE Toulouse) or at MetaGenoPolis (INRAE Jouy-en-Josas) according to the manufacturer’s instructions. Additional details are available in the supplemental material.

The 104 genomes with only Illumina reads were assembled using RiboSeed v0.4.73 (90). RiboSeed uses a reference genome to resolve ribosomal RNA operons and globally improve whole genome assembly. Assemblies were performed using NCTC 12421 (# NZ_LS483306) as reference genome for rRNA operons, using SPAdes v3.13.0 (91) as assembler in “careful” mode using k values of 21, 33, 55, 77 and 99. The 14 genomes with Illumina and Nanopore reads were performed using Unicycler 0.4.4 (92), an assembly pipeline for bacterial genomes that uses SPAdes for short read assembly Miniasm and Racon for long read assemblies and polishing. Unicycler was launched with default parameters.

Genome annotation was conducted with Prokka v1.12 (93). First, coding DNA sequences were identified on contigs longer than 200 bp by Prodigal v2.6.1 (94), which penalized CDSs shorter than 250 bp in order to filter out false positives. CDSs were first annotated (--proteins Prokka parameter) using a protein bank extracted from all the *Enterococcus* complete genomes in RefSeq (4198 genomes retrieved in April, 2020). Annotation from hits with an e-value cut-off of 10-9 and 80% coverage were transferred. CDSs with no hits on this bank were annotated using Prokka default workflow and databanks.

### Pan-genome analysis and phylogenomic tree construction

Pan-genome analysis was performed by comparing 118 *de novo* sequenced genomes of *E. cecorum* and 30 public genomes from NCBI with N50 above 20 kb (January 2021). One reference genome was available on the NCBI nucleotide database as NCTC 12421. Protein clustering was performed by Roary (95) v-3.12.0 with 94% of identity, a “percentage of isolates a gene must be in to be core” of 100%, and the parameter “without split paralogs”. Genes classified as core were genes present in all 148 genomes. Accessory genes were all other genes present in 147 or fewer genomes. The gene accumulation curves were produced with ggplot2 (96) from Roary analysis. The phylogenomic analysis was performed using the *E. cecorum* core-genome. The 1,206 core genes of *E. cecorum* were aligned using a codon aware alignment produced by PRANK (v170427); an unrooted tree was then constructed using the BioNJ algorithm (97) in SeaView (v4.2) (98), using the Jukes and Cantor distance and 1,000 bootstrap replicates. Clades were determined using the Jukes and Cantor distance between aligned core-genes of less than 0.021 and a bootstrap value greater than 75%.

### Acquired antimicrobial resistance genes search

ARGs research was performed using the ResFinder (v2.1) (99) tool and database (2019-04-26) for 90% of gene identity and 60% of coverage. BLASTN was used to detect the *bcr*-like gene (*uppP_2*) and to compare the *vanA* locus with pVEF3 of from *E. faecium* 01_233 (34). Positive hits had at least 90% nucleotide identity and 60% coverage. GenoPlotR (v0.8.11) was used to visualize BLASTN results having > 90% identity (34).

### ICE, GI, and Prophage detection

ICEs and IMEs were detected in the eleven complete genomes and the two large contigs of genome CIRMBP-1320 using ICEscreen (https://icescreen.migale.inrae.fr) and then inspected visually for delineation. Genomic islands corresponded to large insertion of more than 10 CDSs resulting in a synteny break between two genomes. Prophage prediction was performed using the prophage detection tools PHASTER (PHAge Search Tool Enhanced Release) (100) and VIBRANT (Virus Identification By iteRative ANnoTation) (101). Only predicted prophages with prediction scoring ≥100 with PHASTER or VIBRANT were retained and manually inspected to determine the attachment and integration sites in reference to the NCTC12421 genome (Accession NZ_LS483306). PHASTER was further used to identify the most similar phage genomes.

### Hierarchical clustering

Estimated marginal mean of biofilm and adhesion to collagen biovolumes, and serum growth indexes were adjusted for experiment and strain factors using a linear model with the “emmeans” R package (1.4). Strains with a similar Euclidian distance between estimated marginal means were grouped using a hierarchical clustering algorithm with average linkage. Clusters of strains were defined by cutting dynamically the dendrogram, using the DynamicTreeCut R package (1.63-1) (102).

### Genome wide association study (GWAS)

GWAS analyses were performed using TreeWas (1.0) (103) to identify genetic loci (SNP and gene presence/absence) associated with clade membership (clade A, B, C, D or E), clinical origin (clinical and non-clinical poultry isolates and clinical human isolates), biofilm robustness (as binary strong values versus others, as binary weak values versus others and phenotypic groups obtained by clustering), and collagen type II adhesion and/or growth in chicken serum (similarly as biofilm robustness). Significant genetic loci corresponded to p-value less than or equal to 0.05 according to terminal test.

A set of criteria was applied to select SNPs or genes of interest. Criteria applied to select clade-specific SNPs were more stringent and only group-exclusive SNPs were retained (sensitivity = 1 or 0 and specificity = 1 or 0). Genes whose presence was associated with clade membership (clade A, C, or E) or clinical origin (clinical and non-clinical poultry isolates, and clinical human isolates), biofilm robustness, collagen type II adhesion, or growth in chicken serum had sensitivity and specificity scores greater than 0.66. In addition, genes whose absence was associated with clade membership or clinical origin, biofilm robustness, collagen type II adhesion, or growth in chicken serum, had sensitivity and specificity scores below 0.33.

## Data availability

All genomic data have been deposited in the EMBL ENA database under the project number ERP135100. Accession numbers of raw reads and assembled genomes are available in Table S2A.

## Acknowledgements

The authors thank Drs L. Borst (North Carolina State University, USA), B. Dolka (Warsaw University of Life Sciences, Poland), E. Oswald (IRSD, Toulouse, France), J. Van Acker Acker (AZ Sint-Lucas, Laboratory of Clinical microbiology, Ghent, Belgium), M. Vaneechoutte (Ghent University Hospital, Belgium), and P. Warnke (Universitätsmedizin Rostock, Germany) for generously providing strains used in this study. We thank Marie Bernard, Marine Gilles, and Arnaud Marie for technical assistance. We are grateful to Julie Puterflam and Jean-Luc Guerin for fruitful discussions and to Luc Devriese for the complementary information on strain NCTC12421. This work has benefited from the facilities and expertise of the MIMA2 MET–GABI (INRAE, AgroParisTech, 78352 Jouy-en-Josas, France; www6.jouy.inra.fr/mima2). We are grateful to the INRAE MIGALE bioinformatics facility (MIGALE, INRAE, 2020. Migale bioinformatics Facility, doi: 10.15454/1.5572390655343293E12) for providing help and/or computing and/or storage resources. Migale is part of the Institut Français de Bioinformatique (ANR-11-INBS-0013). J.L. was supported by a fellowship from ANSES and INRAE. This work was supported by the INRAE metaprogramme GISA (project CecoType) and by the French Ministry of Agriculture (DGAL) through the program Ecoantibio2 N°2018-180.

## SUPPLEMENTAL MATERIAL

### Supplemental methods

**Table S1:** Metadata of strains sequenced in the study.

**Table S2: A:** *E. cecorum* genomes sequenced in this study and included in the comparative analysis. **B:** Non-redundant *E. cecorum* genomes available in the NCBI Data base included in the comparative analysis.

**Table S3:** Genes differentially distributed between clade E isolates and isolates of other clades.

**Table S4:** A: Genes differentially distributed between clinical and non-clinical poultry isolates or clinical human isolates. **B:** Table S4B: Proposed set of discriminant accessory genes.

**Table S5: A:** Transposon, ICE, IME and genomic islands detected in *E. cecorum* complete or high-quality genomes. **B:** Distribution of ICEs and genomic islands of 12 strains in all available *E. cecorum* genomes.

**Table S6: A:** Prophages predicted in *E. cecorum* draft genomes. **B:** *E. cecorum* draft genomes with no predicted prophages.

**Table S7:** Genes present in highly and intermediate virulent *E. cecorum* isolates in embryonated eggs and absent in the non-virulent isolates.

**Figure S1: Inversion in two genomes compared to reference genome.** A. Alignment of CIRMBP-1261 genome and NCTC 12421. B. Alignment of CIRMBP-1287 and NCTC 12421. Blue lines symbolize inversion in genomes.

**Figure S2: Pairwise genetic distances between the *E. cecorum* genomes.** Color scale goes from green for close genomes to red for distant genomes. Tree clades are framed with respective colors used on the phylogenetic tree. Sub-clades are framed in black.

**Figure S3: Alignment of the predicted capsule locus in the complete genomes.** Arrows are annotated CDSs on both elements. The grayscale shading represents regions of nucleotide sequence identity (100% to 94%) determined by BLASTN analysis. The number of isolates that share each type of locus, among the 117 other genomes in the study, is shown on the right.

**Figure S4: Vancomycin and narasin resistance operons alignment between CIRMBP-1294 and *E. faecium* plasmid, pVEF3.** Blue arrows are annotated CDSs on the both elements. Grey highlights represent 99% or more of identity in BLASTN. Frames indicate vancomycin resistance and narasin resistance operons.

## REFERENCES

1. Devriese LA, Dutta GN, Farrow JAE, Vandekerckhove A, Phillips BA. 1983. *Streptococcus cecorum*, a new species isolated from chickens. Int J Syst Bacteriol 33:772–776.

2. Devriese LA, Hommez J, Wijfels R, Haesebrouck F. 1991. Composition of the enterococcal and streptococcal intestinal flora of poultry. J Appl Bacteriol 71:46–50.

3. Schreier J, Rautenschlein S, Jung A. 2021. Different virulence levels of *Enterococcus cecorum* strains in experimentally infected meat-type chickens. PLoS One 16:e0259904.

4. Dolka B, Golębiewska-Kosakowska M, Krajewski K, Kwieciński P, Nowak T, Zubstarski J, Wilczyński J, Szeleszczuk P. 2017. Occurrence of *Enterococcus* spp. in poultry in Poland based on 2014–2015 data. Med Weter 73:220–224.

5. Souillard R, Allain V, Toux JY, Lecaer V, Lahmar A, Tatone F, Amenna-Bernard A, Le Bouquin S. 2019. Synthèse des pathologies aviaires observées en 2018 par le Réseau National d’Observations Épidémiologiques en Aviculture (RNOEA). Bull Epid Santé Anim 88:1–5.

6. Aziz T, Barnes HJ. 2007. Is spondylitis an emerging disease in broiler breeders? World Poultry 12:44–45.

7. Kense MJ, Landman WJ. 2011. *Enterococcus cecorum* infections in broiler breeders and their offspring: molecular epidemiology. Avian Pathol 40:603–12.

8. Stalker MJ, Brash ML, Weisz A, Ouckama RM, Slavic D. 2010. Arthritis and osteomyelitis associated with *Enterococcus cecorum* infection in broiler and broiler breeder chickens in Ontario, Canada. J Vet Diagn Invest 22:643–5.

9. Wood AM, MacKenzie G, McGiliveray NC, Brown L, Devriese LA, Baele M. 2002. Isolation of *Enterococcus cecorum* from bone lesions in broiler chickens. Vet Rec 150:27.

10. Borst LB, Suyemoto MM, Sarsour AH, Harris MC, Martin MP, Strickland JD, Oviedo EO, Barnes HJ. 2017. Pathogenesis of enterococcal spondylitis caused by *Enterococcus cecorum* in broiler chickens. Vet Pathol 54:61–73.

11. Jung A, Rautenschlein S. 2014. Comprehensive report of an *Enterococcus cecorum* infection in a broiler flock in Northern Germany. BMC Vet Res 10:311.

12. Jung A, Chen LR, Suyemoto MM, Barnes HJ, Borst LB. 2018. A review of *Enterococcus cecorum* infection in poultry. Avian Dis 62:261–271.

13. Braga JFV, Martins NRS, Ecco R. 2018. Vertebral osteomyelitis in broilers: a review. Avian Pathol 20:605–616.

14. Jung A, Metzner M, Ryll M. 2017. Comparison of pathogenic and non-pathogenic *Enterococcus cecorum* strains from different animal species. BMC Microbiol 17:33.

15. Martin LT, Martin MP, Barnes HJ. 2011. Experimental reproduction of enterococcal spondylitis in male broiler breeder chickens. Avian Dis 55:273–8.

16. Grund A, Rautenschlein S, Jung A. 2022. Detection of *Enterococcus cecorum* in the drinking system of broiler chickens and examination of its potential to form biofilms. Europ Poult Sci 86:15.

17. De Herdt P, Defoort P, Van Steelant J, Swam H, Tanghe L, Van Goethem S, Vanrobaeys M. 2008. *Enterococcus cecorum* osteomyelitis and arthritis in broiler chickens. Vlaams Diergeneeskd Tijdschr 78:44–48.

18. Grund A, Rautenschlein S, Jung A. 2020. Tenacity of *Enterococcus cecorum* at different environmental conditions. J Appl Microbiol 130:1494–1507.

19. Borst LB, Suyemoto MM, Scholl EH, Fuller FJ, Barnes HJ. 2015. Comparative genomic analysis identifies divergent genomic features of pathogenic *Enterococcus cecorum* including a type IC CRISPR-Cas system, a capsule locus, an *epa-like* locus, and putative host tissue binding proteins. PLoS One 10:e0121294.

20. Paganelli FL, Leavis HL, He S, van Sorge NM, Payré C, Lambeau G, Willems RJL, Rooijakkers SHM. 2018. Group IIA-secreted phospholipase A(2) in human serum kills commensal but not clinical *Enterococcus faecium* isolates. Infect Immun 86.

21. Borst LB, Suyemoto MM, Keelara S, Dunningan SE, Guy JS, Barnes HJ. 2014. A chicken embryo lethality assay for pathogenic *Enterococcus cecorum*. Avian Dis 58:244–8.

22. Jackson CR, Kariyawasam S, Borst LB, Frye JG, Barrett JB, Hiott LM, Woodley TA. 2015. Antimicrobial resistance, virulence determinants and genetic profiles of clinical and nonclinical *Enterococcus cecorum* from poultry. Lett Appl Microbiol 60:111–9.

23. Boerlin P, Nicholson V, Brash M, Slavic D, Boyen F, Sanei B, Butaye P. 2012. Diversity of *Enterococcus cecorum* from chickens. Vet Microbiol 157:405–11.

24. Borst LB, Suyemoto MM, Robbins KM, Lyman RL, Martin MP, Barnes HJ. 2012. Molecular epidemiology of *Enterococcus cecorum* isolates recovered from enterococcal spondylitis outbreaks in the southeastern United States. Avian Pathol 41:479–85.

25. Dolka B, Chrobak-Chmiel D, Makrai L, Szeleszczuk P. 2016. Phenotypic and genotypic characterization of *Enterococcus cecorum* strains associated with infections in poultry. BMC Vet Res 12:129.

26. Robbins KM, Suyemoto MM, Lyman RL, Martin MP, Barnes HJ, Borst LB. 2012. An outbreak and source investigation of enterococcal spondylitis in broilers caused by *Enterococcus cecorum*. Avian Dis 56:768–73.

27. Wijetunge DS, Dunn P, Wallner-Pendleton E, Lintner V, Lu H, Kariyawasam S. 2012. Fingerprinting of poultry isolates of *Enterococcus cecorum* using three molecular typing methods. J Vet Diagn Invest 24:1166–71.

28. Sharma P, Gupta SK, Barrett JB, Hiott LM, Woodley TA, Kariyawasam S, Frye JG, Jackson CR. 2020. Comparison of antimicrobial resistance and pan-genome of clinical and non-clinical *Enterococcus cecorum* from poultry using whole-genome sequencing. Foods 9:686.

29. Dolka B, Boyen F, Butaye P, Heidemann Olsen R, Naundrup Thofner IC, Christensen JP. 2015. Draft genome sequences of two commensal *Enterococcus cecorum* strains isolated from chickens in Belgium. Genome Announc 3.

30. Dolka B, Heidemann Olsen R, Naundrup Thofner IC, Christensen JP. 2015. Draft genome sequences of five clinical *Enterococcus cecorum* strains isolated from different poultry species in Poland. Genome Announc 3.

31. Dutta GN, Devriese LA. 1982. Susceptibility of fecal streptococci of poultry origin to nine growth-promoting agents. Appl Environ Microbiol 44:832–7.

32. Yew WS, Gerlt JA. 2002. Utilization of L-ascorbate by *Escherichia coli* K-12: assignments of functions to products of the *yjf-sga* and *yia-sgb* operons. J Bacteriol 184:302–6.

33. Linares D, Michaud P, Delort AM, Traikia M, Warrand J. 2011. Catabolism of L-ascorbate by *Lactobacillus rhamnosus* GG. J Agric Food Chem 59:4140–7.

34. Nilsson O, Myrenas M, Agren J. 2016. Transferable genes putatively conferring elevated minimum inhibitory concentrations of narasin in *Enterococcus faecium* from Swedish broilers. Vet Microbiol 184:80–3.

35. Nilsson O, Greko C, Bengtsson B, Englund S. 2012. Genetic diversity among VRE isolates from Swedish broilers with the coincidental finding of transferrable decreased susceptibility to narasin. J Appl Microbiol 112:716–22.

36. Bellanger X, Payot S, Leblond-Bourget N, Guedon G. 2014. Conjugative and mobilizable genomic islands in bacteria: evolution and diversity. FEMS Microbiol Rev 38:720–60.

37. Ambroset C, Coluzzi C, Guedon G, Devignes MD, Loux V, Lacroix T, Payot S, Leblond-Bourget N. 2015. New insights into the classification and integration specificity of *Streptococcus* integrative conjugative elements through extensive genome exploration. Front Microbiol 6:1483.

38. Coluzzi C, Guédon G, Devignes M-D, Ambroset C, Loux V, Lacroix T, Payot S, Leblond-Bourget N. 2017. A glimpse into the world of integrative and mobilizable elements in streptococci reveals an unexpected diversity and novel families of mobilization proteins. Front Microbiol 8:443.

39. Cornuault JK, Petit MA, Mariadassou M, Benevides L, Moncaut E, Langella P, Sokol H, De Paepe M. 2018. Phages infecting *Faecalibacterium prausnitzii* belong to novel viral genera that help to decipher intestinal viromes. Microbiome 6:65.

40. Yoon BH, Chang HI. 2015. Genomic annotation for the temperate phage EFC-1, isolated from *Enterococcus faecalis* KBL101. Arch Virol 160:601–4.

41. Díaz E, López R, García JL. 1992. EJ-1, a temperate bacteriophage of *Streptococcus pneumoniae* with a Myoviridae morphotype. J Bacteriol 174:5516–25.

42. Mills S, Griffin C, O’Sullivan O, Coffey A, McAuliffe OE, Meijer WC, Serrano LM, Ross RP. 2011. A new phage on the ‘Mozzarella’ block: Bacteriophage 5093 shares a low level of homology with other *Streptococcus thermophilus* phages. Int Dairy J 21:963–969.

43. Tettelin H, Masignani V, Cieslewicz MJ, Donati C, Medini D, Ward NL, Angiuoli SV, Crabtree J, Jones AL, Durkin AS, Deboy RT, Davidsen TM, Mora M, Scarselli M, Margarit y Ros I, Peterson JD, Hauser CR, Sundaram JP, Nelson WC, Madupu R, Brinkac LM, Dodson RJ, Rosovitz MJ, Sullivan SA, Daugherty SC, Haft DH, Selengut J, Gwinn ML, Zhou L, Zafar N, Khouri H, Radune D, Dimitrov G, Watkins K, O’Connor KJ, Smith S, Utterback TR, White O, Rubens CE, Grandi G, Madoff LC, Kasper DL, Telford JL, Wessels MR, Rappuoli R, Fraser CM. 2005. Genome analysis of multiple pathogenic isolates of *Streptococcus agalactiae:* implications for the microbial “pan-genome”. Proc Natl Acad Sci USA 102:13950–5.

44. McInerney JO, McNally A, O’Connell MJ. 2017. Why prokaryotes have pangenomes. Nat Microbiol 2:17040.

45. Land M, Hauser L, Jun SR, Nookaew I, Leuze MR, Ahn TH, Karpinets T, Lund O, Kora G, Wassenaar T, Poudel S, Ussery DW. 2015. Insights from 20 years of bacterial genome sequencing. Funct Integr Genomics 15:141–61.

46. Touchon M, Perrin A, de Sousa JAM, Vangchhia B, Burn S, O’Brien CL, Denamur E, Gordon D, Rocha EP. 2020. Phylogenetic background and habitat drive the genetic diversification of *Escherichia coli*. PLoS Genet 16:e1008866.

47. Park CJ, Andam CP. 2020. Distinct but intertwined evolutionary histories of multiple *Salmonella enterica* subspecies. mSystems 5.

48. Weinert LA, Chaudhuri RR, Wang J, Peters SE, Corander J, Jombart T, Baig A, Howell KJ, Vehkala M, Välimäki N, Harris D, Chieu TT, Van Vinh Chau N, Campbell J, Schultsz C, Parkhill J, Bentley SD, Langford PR, Rycroft AN, Wren BW, Farrar J, Baker S, Hoa NT, Holden MT, Tucker AW, Maskell DJ. 2015. Genomic signatures of human and animal disease in the zoonotic pathogen *Streptococcus suis*. Nat Commun 6:6740.

49. Weinert LA, Welch JJ. 2017. Why might bacterial pathogens have small genomes? Trends Ecol Evol 32:936–947.

50. Rosinski-Chupin I, Sauvage E, Mairey B, Mangenot S, Ma L, Da Cunha V, Rusniok C, Bouchier C, Barbe V, Glaser P. 2013. Reductive evolution in *Streptococcus agalactiae* and the emergence of a host adapted lineage. BMC Genomics 14:252.

51. Merhej V, Georgiades K, Raoult D. 2013. Postgenomic analysis of bacterial pathogens repertoire reveals genome reduction rather than virulence factors. Brief Funct Genomics 12:291–304.

52. Souillard R, Laurentie J, Kempf I, Le Caer V, Le Bouquin S, Serror P, Allain V. 2022. Increasing incidence of *Enterococcus*-associated diseases in poultry in France over the past 15 years. Vet Microbiol 269:109426.

53. Hiemstra SJ, Ten Napel J. 2013. Study of the impact of genetic selection on the welfare of chickens bred and kept for meat production (DG SANCO/2011/12254). J Final report of a project commissioned by the European Commission.

54. Delaunay E, Abat C, Rolain JM. 2015. *Enterococcus cecorum* human infection, France. New Microbes New Infect 7:50–1.

55. Stubljar D, Skvarc M. 2015. *Enterococcus cecorum* infection in two critically ill children and in two adult septic patients. Slov Vet Res 52:39–44.

56. Brückner C, Straube E, Petersen I, Sachse S, Keller P, Layher F, Matziolis G, Spiegl U, Zajonz D, Edel M, Roth A. 2019. Low-grade infections as a possible cause of arthrofibrosis after total knee arthroplasty. Patient Saf Surg 13:1.

57. Lebreton F, Manson AL, Saavedra JT, Straub TJ, Earl AM, Gilmore MS. 2017. Tracing the enterococci from paleozoic origins to the hospital. Cell 169:849–861.e13.

58. Khan A, Davlieva M, Panesso D, Rincon S, Miller WR, Diaz L, Reyes J, Cruz MR, Pemberton O, Nguyen AH, Siegel SD, Planet PJ, Narechania A, Latorre M, Rios R, Singh KV, Ton-That H, Garsin DA, Tran TT, Shamoo Y, Arias CA. 2019. Antimicrobial sensing coupled with cell membrane remodeling mediates antibiotic resistance and virulence in *Enterococcus faecalis*. Proc Natl Acad Sci USA 116:26925–32.

59. Frumkin I, Lajoie MJ, Gregg CJ, Hornung G, Church GM, Pilpel Y. 2018. Codon usage of highly expressed genes affects proteome-wide translation efficiency. Proc Natl Acad Sci USA 115:E4940–E4949.

60. Ramos Y, Sansone S, Morales DK. 2021. Sugarcoating it: Enterococcal polysaccharides as key modulators of host-pathogen interactions. PLoS Pathog 17:e1009822.

61. Thurlow LR, Thomas VC, Fleming SD, Hancock LE. 2009. *Enterococcus faecalis* capsular polysaccharide serotypes C and D and their contributions to host innate immune evasion. Infect Immun 77:5551–7.

62. Rohmer L, Hocquet D, Miller SI. 2011. Are pathogenic bacteria just looking for food? Metabolism and microbial pathogenesis. Trends Microbiol 19:341–8.

63. Nogales J, Garmendia J. 2022. Bacterial metabolism and pathogenesis intimate intertwining: time for metabolic modelling to come into action. Microb Biotechnol 15:95–102.

64. Cury J, Touchon M, Rocha EPC. 2017. Integrative and conjugative elements and their hosts: composition, distribution and organization. Nucleic Acids Res 45:8943–8956.

65. Lao J, Guedon G, Lacroix T, Charron-Bourgoin F, Libante V, Loux V, Chiapello H, Payot S, Leblond-Bourget N. 2020. Abundance, Diversity and Role of ICEs and IMEs in the Adaptation of *Streptococcus salivarius* to the Environment. Genes (Basel) 11.

66. Couturier E, Rocha EP. 2006. Replication-associated gene dosage effects shape the genomes of fast-growing bacteria but only for transcription and translation genes. Mol Microbiol 59:1506–18.

67. Lato DF, Golding GB. 2020. Spatial patterns of gene expression in bacterial genomes. J Mol Evol 88:510–520.

68. Goessweiner-Mohr N, Arends K, Keller W, Grohmann E. 2014. Conjugation in Gram-Positive bacteria. Microbiol Spectr 2:Plas-0004-2013.

69. Bhatty M, Laverde Gomez JA, Christie PJ. 2013. The expanding bacterial type IV secretion lexicon. Res Microbiol 164:620–39.

70. Deng L, Spencer BL, Holmes JA, Mu R, Rego S, Weston TA, Hu Y, Sanches GF, Yoon S, Park N, Nagao PE, Jenkinson HF, Thornton JA, Seo KS, Nobbs AH, Doran KS. 2019. The Group B Streptococcal surface antigen I/II protein, BspC, interacts with host vimentin to promote adherence to brain endothelium and inflammation during the pathogenesis of meningitis. PLoS Pathog 15:e1007848.

71. Chuzeville S, Dramsi S, Madec JY, Haenni M, Payot S. 2015. Antigen I/II encoded by integrative and conjugative elements of *Streptococcus agalactiae* and role in biofilm formation. Microb Pathog 88:1–9.

72. Jurėnas D, Fraikin N, Goormaghtigh F, Van Melderen L. 2022. Biology and evolution of bacterial toxin-antitoxin systems. Nat Rev Microbiol 20:335–350.

73. Couvigny B, Lapaque N, Rigottier-Gois L, Guillot A, Chat S, Meylheuc T, Kulakauskas S, Rohde M, Mistou MY, Renault P, Dore J, Briandet R, Serror P, Guedon E. 2017. Three glycosylated serine-rich repeat proteins play a pivotal role in adhesion and colonization of the pioneer commensal bacterium, *Streptococcus salivarius*. Environ Microbiol 19:3579–3594.

74. Latousakis D, MacKenzie DA, Telatin A, Juge N. 2020. Serine-rich repeat proteins from gut microbes. Gut Microbes 11:102–117.

75. Kaufman JH, Terrizzano I, Nayar G, Seabolt E, Agarwal A, Slizovskiy IB, Noyes N. 2020. Integrative and Conjugative Elements (ICE) and Associated Cargo Genes within and across Hundreds of Bacterial Genera. bioRxiv https://doi.org/10.1101/2020.04.07.030320:2020.04.07.030320.

76. Roberts MC, Schwarz S. 2016. Tetracycline and phenicol resistance genes and mechanisms: importance for agriculture, the environment, and humans. J Environ Qual 45:576–92.

77. Lyras D, Adams V, Ballard SA, Teng WL, Howarth PM, Crellin PK, Bannam TL, Songer JG, Rood JI. 2009. tIS*Cpe8*, an IS *1595*-family lincomycin resistance element located on a conjugative plasmid in *Clostridium perfringens*. J Bacteriol 191:6345–51.

78. Han X, Du XD, Southey L, Bulach DM, Seemann T, Yan XX, Bannam TL, Rood JI. 2015. Functional analysis of a bacitracin resistance determinant located on ICE*Cp1*, a novel Tn*916*-like element from a conjugative plasmid in *Clostridium perfringens*. Antimicrob Agents Chemother 59:6855–65.

79. Chen M-Y, Lira F, Liang H-Q, Wu R-T, Duan J-H, Liao X-P, Martínez JL, Liu Y-H, Sun J. 2016. Multilevel selection of *bcrABDR*-mediated bacitracin resistance in *Enterococcus faecalis* from chicken farms. Sci Rep 6:1–7.

80. Reynolds LJ, Anjum MF, Roberts AP. 2020. Detection of a novel, and likely ancestral, Tn*916*-like element from a human saliva metagenomic library. Genes 11:548.

81. de Vries LE, Vallès Y, Agersø Y, Vaishampayan PA, García-Montaner A, Kuehl JV, Christensen H, Barlow M, Francino MP. 2011. The gut as reservoir of antibiotic resistance: microbial diversity of tetracycline resistance in mother and infant. PLOS ONE 6:e21644.

82. Harada T, Kawahara R, Kanki M, Taguchi M, Kumeda Y. 2012. Isolation and characterization of *vanA* genotype vancomycin-resistant *Enterococcus cecorum* from retail poultry in Japan. Int J Food Microbiol 153:372–7.

83. Singer RS, Porter LJ, Schrag NFD, Davies PR, Apley MD, Bjork K. 2020. Estimates of on-farm antimicrobial usage in broiler chicken production in the United States, 2013-2017. Zoonoses Public Health 67 Suppl 1:22–35.

84. Debaere O. 2016. Ecoantibio: premier plan de réduction des risques d’antibiorésistance en médecine vétérinaire (2012–2016). Bull Acad Vet Fr 169:186–189.

85. Jouvin-Marche E, Carrara G, Pulcini C, Andremont A, Danan C, Couderc-Obert C, Lienhardt C, Kieny M-P, Yazdanpanah YJTL. 2020. French research strategy to tackle antimicrobial resistance. Lancet 395:1239–1241.

86. Urban D, Chevance A, Moulin G. 2021. Surveillance des ventes de médicaments vétérinaires contenant des antibiotiques en France en 2020: Rapport annuel. Anses, Agence nationale de sécurité sanitaire de l’alimentation, de l’environnement et du travail 1–92 doi:https://hal-anses.archives-ouvertes.fr/anses-03515142.

87. Monteiro G, Rossi D, Valadares Jr E, Peres P, Braz R, Notário F, Gomes M, Silva R, Carrijo K, Fonseca B. 2021. Lactic bacterium and *Bacillus* Sp. biofilms can decrease the viability of *Salmonella gallinarum*, *Salmonella heidelberg*, *Campylobacter jejuni* and methicillin resistant *Staphylococcus aureus* on different substrates. Braz J Poultry Sci 23.

88. Wideman RF, Al-Rubaye A, Kwon YM, Blankenship J, Lester H, Mitchell KN, Pevzner IY, Lohrmann T, Schleifer J. 2015. Prophylactic administration of a combined prebiotic and probiotic, or therapeutic administration of enrofloxacin, to reduce the incidence of bacterial chondronecrosis with osteomyelitis in broilers. Poult Sci 94:25–36.

89. Luise D, Bosi P, Raff L, Amatucci L, Virdis S, Trevisi P. 2022. *Bacillus* spp. probiotic strains as a potential tool for limiting the use of antibiotics, and improving the growth and health of pigs and chickens. Front Microbiol 13.

90. Waters NR, Abram F, Brennan F, Holmes A, Pritchard L. 2018. riboSeed: leveraging prokaryotic genomic architecture to assemble across ribosomal regions. Nucleic Acids Res 46:e68.

91. Bankevich A, Nurk S, Antipov D, Gurevich AA, Dvorkin M, Kulikov AS, Lesin VM, Nikolenko SI, Pham S, Prjibelski AD, Pyshkin AV, Sirotkin AV, Vyahhi N, Tesler G, Alekseyev MA, Pevzner PA. 2012. SPAdes: a new genome assembly algorithm and its applications to single-cell sequencing. J Comput Biol 19:455–77.

92. Wick RR, Judd LM, Gorrie CL, Holt KE. 2017. Unicycler: Resolving bacterial genome assemblies from short and long sequencing reads. PLoS Comput Biol 13:e1005595.

93. Ewels P, Magnusson M, Lundin S, Käller M. 2016. MultiQC: summarize analysis results for multiple tools and samples in a single report. Bioinformatics 32:3047–8.

94. Hyatt D, Chen GL, Locascio PF, Land ML, Larimer FW, Hauser LJ. 2010. Prodigal: prokaryotic gene recognition and translation initiation site identification. BMC Bioinformatics 11:119.

95. Page AJ, Cummins CA, Hunt M, Wong VK, Reuter S, Holden MT, Fookes M, Falush D, Keane JA, Parkhill J. 2015. Roary: rapid large-scale prokaryote pan genome analysis. Bioinformatics 31:3691–3.

96. Wickham H. 2016. ggplot2: Elegant Graphics for Data Analysis. Springer-Verlag New York.

97. Gascuel O. 1997. BIONJ: an improved version of the NJ algorithm based on a simple model of sequence data. Mol Biol Evol 14:685–95.

98. Gouy M, Guindon S, Gascuel O, evolution. 2010. SeaView version 4: a multiplatform graphical user interface for sequence alignment and phylogenetic tree building. J Molecular biology 27:221–224.

99. Zankari E, Hasman H, Cosentino S, Vestergaard M, Rasmussen S, Lund O, Aarestrup FM, Larsen MV. 2012. Identification of acquired antimicrobial resistance genes. J Antimicrob Chemother 67:2640–4.

100. Arndt D, Grant JR, Marcu A, Sajed T, Pon A, Liang Y, Wishart DS. 2016. PHASTER: a better, faster version of the PHAST phage search tool. Nucleic Acids Res 44:W16–21.

101. Kieft K, Zhou Z, Anantharaman K. 2020. VIBRANT: automated recovery, annotation and curation of microbial viruses, and evaluation of viral community function from genomic sequences. Microbiome 8:90.

102. Langfelder P, Zhang B, Horvath S. 2008. Defining clusters from a hierarchical cluster tree: the Dynamic Tree Cut package for R. J Bioinformatics 24:719–720.

103. Collins C, Didelot X. 2018. A phylogenetic method to perform genome-wide association studies in microbes that accounts for population structure and recombination. PLoS Comput Biol 14:e1005958.

